# A molecular calcium integrator reveals a striatal cell-type driving aversion

**DOI:** 10.1101/2020.11.01.364174

**Authors:** Christina K. Kim, Mateo I. Sanchez, Paul Hoerbelt, Lief E. Fenno, Robert C. Malenka, Karl Deisseroth, Alice Y. Ting

**Affiliations:** Department of Genetics, Stanford University, Stanford, CA 94305, USA; Chan Zuckerberg Biohub, San Francisco, CA 94158, USA; Department of Psychiatry and Behavioral Sciences, Stanford University, Stanford, CA 94305, USA; Department of Bioengineering, Stanford University, Stanford, CA 94035, USA; Howard Hughes Medical Institute, Stanford University, Stanford, CA 94305, USA; Department of Biology, Stanford University, Stanford, CA 94305, USA

**Author notes:** These authors contributed equally to this work (Christina K. Kim, Mateo I. Sanchez).

## Abstract

The ability to record transient cellular events in the DNA or RNA of cells would enable precise, large-scale analysis, selection, and reprogramming of heterogeneous cell populations. Here we report a molecular technology for stable genetic tagging of cells that exhibit activity-related increases in intracellular calcium concentration (FLiCRE). We used FLiCRE to transcriptionally label activated neural ensembles in the nucleus accumbens of the mouse brain during brief stimulation of aversive inputs. Using single-cell RNA sequencing, we detected FLiCRE transcripts among the endogenous transcriptome, providing simultaneous readout of both cell-type and calcium activation history. We identified a cell-type in the nucleus accumbens activated downstream of long-range excitatory projections. Taking advantage of FLiCRE’s modular design, we expressed an optogenetic channel selectively in this cell-type, and showed that direct recruitment of this otherwise genetically-inaccessible population elicits behavioral aversion. The specificity and minute-resolution of FLiCRE enables molecularly-informed characterization, manipulation, and reprogramming of activated cellular ensembles.

## INTRODUCTION

Networks of interconnected cells give rise to tissue and organ function in living systems. For example, in the mammalian brain, functionally coupled neural networks coordinate their spiking activity to process stimuli and give rise to specific behaviors. However, the complexity of many functional cell networks makes them difficult to delineate and characterize. There are few tools available to answer fundamental questions, such as what interconnected cell types comprise a given network, and how the activity of these cells drives behavioral output.

Imaging is frequently used to identify cell subpopulations with activity correlated to a particular behavior, experience, or cognition. In the brain, real-time imaging of fluorescent activity sensors can detect neurons that exhibit time-locked spiking activity with brief stimuli such as visual input (Chen et al., 2013; Gong et al., 2015), or neurons that elevate their activity during more complex behaviors such as changes in motivation (Burnett et al., 2016; Calipari et al., 2016). However, while real-time imaging can record activity dynamics with sub-second timescale precision, it is limited to relatively small fields of view and requires custom-built optical instrumentation to directly combine it with other methods of characterization, such as transcriptome sequencing (Lee et al., 2019) or activity-guided optogenetic manipulations during behavior (Carrillo-Reid et al., 2019; Jennings et al., 2019; Marshel et al., 2019).

To overcome the inherent limitations of imaging, several technologies have been developed for stable, activity-dependent tagging of cell populations. TRAP (Allen et al., 2017; Guenthner et al., 2013), ESARE (Kawashima et al., 2013; Kim et al., 2017) and tetTag (Liu et al., 2012; Reijmers et al., 2007) give fluorescent transgene expression in neuron subpopulations that turn on immediate early gene (IEG) expression (a marker of neuronal activity in mammalian systems) during a drug-delivery window of ~6 hrs (DeNardo et al., 2019). Subsequent imaging of the transgene can reveal the cells that were previously active and some of their characteristics, such as anatomical location and projection targets (DeNardo et al., 2019; Ye et al., 2016). Immunoprecipitation or FACS enrichment of the IEG-driven transgene can also allow transcriptome sequencing of activated cells; moreover, if the selected transgene is a microbial opsin (Deisseroth, 2015) such as a channelrhodopsin (Boyden et al., 2005) or other molecular actuator, then optogenetic manipulation can reveal the causal relationship between tagged neurons and behavior (Allen et al., 2017; Ye et al., 2016). These tools, however, have two major limitations. First, the tagging time window is far longer than the timescale of most behaviors of interest, due to the both the slow onset of IEG promoter activity, and the time it takes for the gating drug to be metabolized (e.g., tamoxifen, used to turn on IEG-driven Cre-ER). This results in a degradation of specificity and insufficient signal-to-noise for tagging neurons active during short-lived behaviors. Second, the precise relationship between IEG expression and spiking activity in neurons has not been broadly characterized and varies substantially among neuronal sub-types in different brain regions (Kawashima et al., 2014).

A more temporally-specific and broadly applicable technology would provide activity-dependent cell tagging based on calcium signals rather than IEG expression. Increases in intracellular calcium faithfully track action potentials on the timescale of milliseconds instead of hours (Lev-Ram and Grinvald, 1987). A recent innovative technology, CaMPARI (Fosque et al., 2015), is a photoswitchable fluorescent protein that turns from green to red in the presence of both ultraviolet light and high calcium; however, CaMPARI does not have the versatility of transcription-based tools; for example, it cannot be used to drive selective channelrhodopsin expression for subsequent manipulation of tagged cell subpopulations.

To further enable the study of functional cell networks – their activity, molecular characteristics, and causal relationship to behavior – we envisioned a “calcium molecular integrator” that can integrate transient increases in intracellular calcium, and then store a stable memory of this calcium signal as a new, modular transcript expressed in the cell. This recorder should have sufficient sensitivity and temporal resolution to detect calcium signals even within brief user-specified time windows, on the timescale of minutes instead of hours. By storing calcium signal history in the form of a new transcript, the activity can be directly read out using high-throughput single-cell RNA sequencing technology. This massively parallel strategy reports the entire endogenous transcriptome, and thus cell-type identity, of each non-active and active cell during the time window of recording. In addition, the new transcript can be an actuator that enables direct manipulation of the previously active cells *in vivo,* linking the cell-type identity discovered through sequencing with functional effects on behavior.

Here we report FLiCRE (Fast Light and Calcium-Regulated Expression), an engineered tool that enables molecular and functional characterization of acutely activated cell ensembles with minute-timescale sensitivity in mice. We apply FLiCRE to label neurons in the nucleus accumbens (NAc) that are activated during brief optogenetic stimulation of long-range excitatory inputs thought to modulate motivated behaviors. By profiling these activated FLiCRE neurons using single-cell RNA sequencing, we identify unique molecular features of a distinct NAc celltype coupled to these excitatory inputs. We then genetically control this unique cell-type using a FLiCRE-driven channelrhodopsin to discover a functional role in driving behavioral aversion.

## RESULTS

For the design of FLiCRE, we envisioned a transcription factor with activity gated by both calcium and blue light (“AND” logic). Calcium is a near-universal activation signal, not only in neurons but also in many other cell types, such as T cells in the immune system upon activation (Le Borgne et al., 2016) and cardiomyocytes during contraction (Baylor and Hollingworth, 2011). We selected light as our temporal gate, because in contrast to drugs such as tamoxifen, light can be delivered to relevant tissue and shut off rapidly. The use of a transcription factor provides versatile genetic access to the activated cells, enabling expression of any transgene of interest. The transgene can be read out by imaging or sequencing, or drive expression of an actuator for manipulation.

The design of FLiCRE is shown in **Figure 1A**. FLiCRE tethers a transcription factor to the plasma membrane via a transmembrane domain, the calmodulin-binding peptide MKII, an evolved light-oxygen-voltage (eLOV) photosensory domain, and a tobacco etch virus cleavage site (TEVcs). A separate construct – truncated TEV protease (TEVp) fused to calcium-sensing calmodulin – is expressed in the cytosol at low levels. In the presence of high intracellular calcium, TEVp-calmodulin interacts with MKII, bringing the protease into proximity of the TEVcs. However, in the absence of blue light to activate LOV, the TEVcs remains sterically caged and inaccessible. When both blue light *and* high intracellular calcium are present, the TEVp cleaves the TEVcs, releasing the transcription factor from the membrane to translocate to the nucleus and drive transgene expression. FLiCRE’s design is informed by our previous tool, FLARE (Wang et al., 2017), but several key design features improve its performance and enable application *in vivo.*

**Figure 1.**
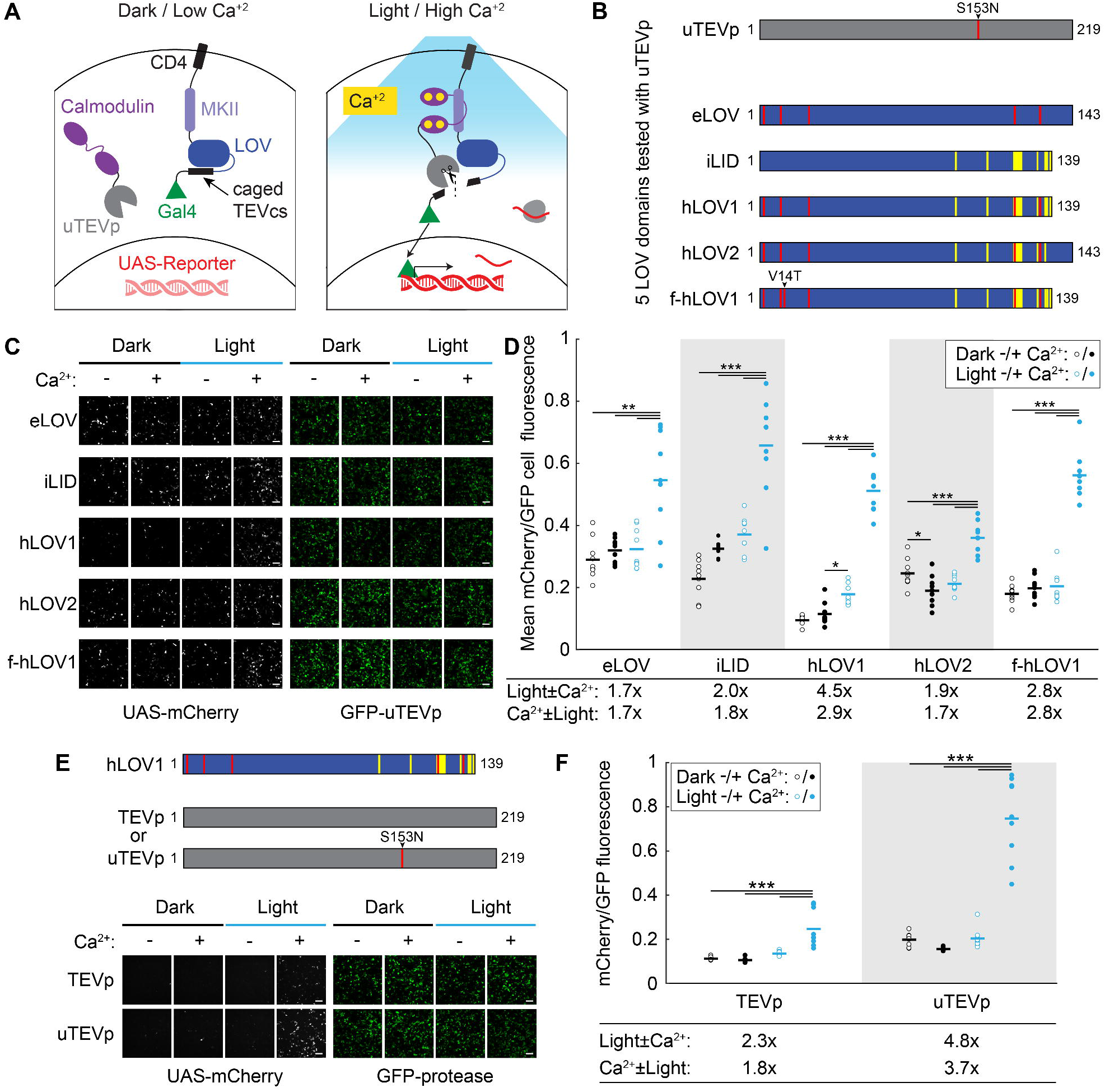
Design and characterization of FLiCRE in cultured HEK293T cells. **(A)** Schematic of FLiCRE. In the basal state (dark/low Ca^2+^), a non-native transcription factor (Gal4) is tethered to the cell membrane via CD4, MKII, LOV, and TEVcs (TEV cleavage site). LOV cages the TEVcs in its inactivated dark state. An evolved TEV protease (uTEVp) is fused to calmodulin (CaM) and expressed cytosolically. In the presence of high Ca^2+^, CaM and MKII interact, bringing uTEVp into proximity of TEVcs, which is only accessible for cleavage when LOV is activated by blue light. Thus both high intracellular Ca^2+^ *and* blue light are required for Gal4 release and translocation to the nucleus to drive expression of a UAS-reporter. **(B)** Optimization of FLiCRE’s LOV domain. Construct designs of five different LOV domains tested with uTEVp (S153N mutation) in FLiCRE. **(C)** FLiCRE’s performance in HEK293T cells transfected with GFP-CaM-uTEVp, UAS-mCherry, and a transmembrane CD4-MKII-LOV-TEVcs-Gal4 component where LOV is one of the 5 LOV variants in **panel B**. Example fluorescence images of UAS-mCherry activation expression (left) and GFP-CaM-uTEVp expression (right), taken ~8 hrs after 10 min of treatment with Ca^2+^ and blue light, alongside control conditions. Scale bars, 100 μm. **(D)** Quantification of experiment in **panel C**. Data represent the mCherry/GFP fluorescence intensity ratio averaged across all cells in a FOV. For all LOV variants, there was a higher mCherry/GFP ratio in the light + Ca^2+^ condition compared to control conditions (*N* = 9 FOVs per condition, 2-way ANOVA interaction F_(1,32)_^eLOV^=8.97, F_(1,32)_^iLID^=9.87, F_(1,32)_^hLOV1^=115.35, F_(1,32)_^hLOV2^=48.84, F_(1,32)_^f-hLOV1^=99.74, *P* < 0.01; Tukey’s multiple comparison’s test, \*\*\**P* < 0.001, \**P* < 0.05). **(E)** Comparison of FLiCRE performance in HEK293T cells transfected with CD4-MKII-hLOV1-TEVcs-Gal4, UAS-mCherry, and either GFP-CaM-TEVp or -uTEVp. Fluorescence images of UAS-mCherry activation and GFP-CaM-TEVp/uTEVp expression following 5 min of Ca^2+^ and blue light. Scale bars, 100 μm. **(F)** Quantification of experiment in **panel E**. For both TEVp and uTEVp, there was a higher mCherry/GFP fluorescence ratio in the light + Ca^2+^ condition compared to control conditions (*N* = 9 FOVs per condition, 2-way ANOVA interaction F_(1,32)_^TEVp^=17.41, F_(1,32)_^uTEVp^=84.73, *P* < 0.001; Tukey’s multiple comparison’s test, \*\*\**P* < 0.0001). **See also Figure S1.**

First, tagging with FLARE required 5 min of continuous calcium and light stimulation in cultured HEK293T cells, and 15 min in cultured neurons; this was due primarily to the slow catalytic activity of the TEV protease (0.4 turnovers/sec). We recently reported ultra TEVp (uTEVp; (Sanchez and Ting, 2020)), a TEVp mutant with >5-fold faster turnover compared to the original truncated TEVp used in FLARE. To generate FLiCRE we first replaced TEVp with uTEVp and tested its performance in HEK293T cells stimulated with blue light and calcium. **Figures 1B and 1C** shows that FLiCRE gives higher UAS-mCherry signal than FLARE, but the higher efficiency of uTEVp also increased background reporter expression in the absence of blue light. This suggests that eLOV was not sufficiently caging the TEVcs in the dark state. Thus we performed rational engineering of eLOV, incorporating mutations and truncations of a previously published LOV domain, iLID (Guntas et al., 2015). In total, we tested 5 different LOV domains with uTEVp (**Figures 1B and 1C**). We found that a truncated version of eLOV that incorporated 9 additional mutations, termed hLOV1, provided the best signal-to-noise of UAS-mCherry FLiCRE reporter expression (**Figure 1D**). hLOV1 also had faster kinetics compared to eLOV, and the addition of the V14T mutation (to give f-hLOV1; V416T in (Kawano et al., 2013)) further improved the kinetics while maintaining sufficient caging of the TEVcs in the dark (**Figures S1A and S1B**). With the incorporation of both uTEVp and the tighter-caging hLOV1, we could achieve over 100% increase in signal-to-noise of UAS-mCherry reporter expression following light and calcium delivery in HEK 293T cells, compared to when using the original FLARE (**Figures 1E and 1F**).

We then tested the capabilities of FLiCRE in cultured cortical neurons using tTA as the transcription factor and TRE-mCherry as the FLiCRE reporter gene. We expressed the FLiCRE constructs in neurons via adeno-associated virus (AAV) infection. To elicit action potentials and increase intracellular calcium, we delivered a varying number of electric field stimulation (E-stim) pulses (4, 8, 16, or 32 pulses delivered at 20Hz every 3 s) to neurons along with blue light for 1 min. Intracellular patchclamp recordings confirmed that our stimulation pulses reliably elicited single action potentials in neurons with 100% fidelity (**Figure 2A**). We found that FLiCRE required between 144 and 256 total E-stim pulses (inferred as action potentials) over 1 min to result in robust labeling (**Figures 2B and 2C**). While keeping the number of pulses per train constant but increasing the total recording time, we also observed increasing amounts of TRE-mCherry expression (**Figures 2D and 2E**). Under these conditions, we could observe significant FLiCRE TRE-mCherry expression within 30 s (a total of 208 pulses). When decreasing the frequency of pulse train delivery to every 6s instead of every 3s, we also observed an overall reduction in TRE-mCherry expression (**Figures S1C and S1D**). Across representative labeling conditions in cultured neurons, FLiCRE’s signal-to-noise ratios ranged from 11x to 66x for light ± E-stim, and 23x to 110x for E-stim ± light (**Figures S1E–S1G**).

**Figure 2.**
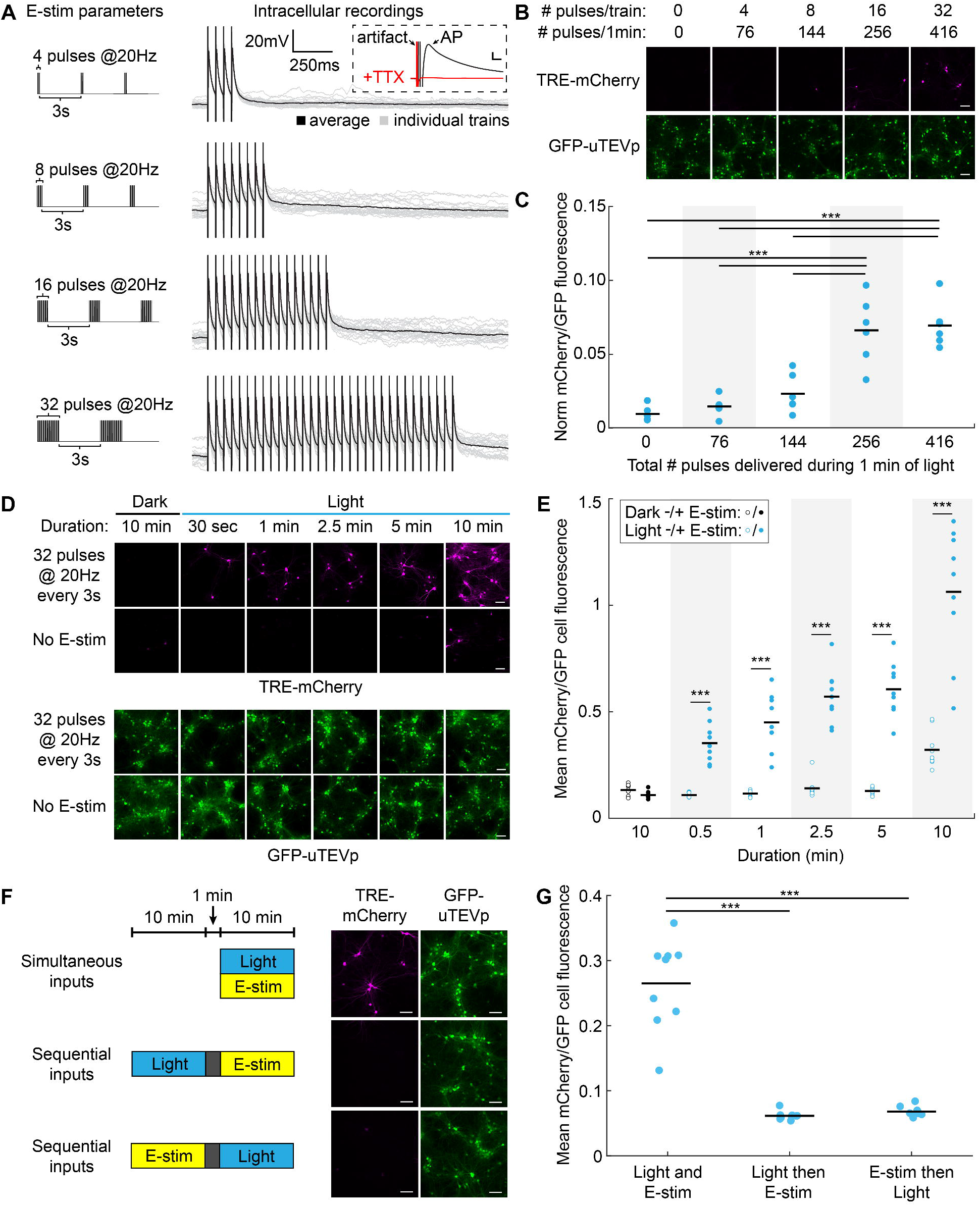
Characterization of FLiCRE’s sensitivity and reversibility in cultured neurons. **(A)** Left: Electric field stimulation (E-stim) parameters used to elicit action potentials during FLiCRE recordings. Right: Intracellular patchclamp recordings in uninfected neurons showing that E-stim pulses elicit single action potentials (100% fidelity in spiking across *N* = 4 cells and 20 trials each). Inset: action potentials were blocked with TTX. Scale bars, 10mV and 5ms. **(B)** Relationship between action potential firing and FLiCRE activation. For FLiCRE expression in neurons, CD4 was replaced with a truncated Neurexin3B fragment, Gal4 replaced with tTA, and a TRE-mCherry reporter gene was used. Fluorescence images of TRE-mCherry activation and GFP-CaM-uTEVp expression were taken ~18 hrs following 1min of light and E-stim (same parameters as in **panel A**). Scale bars, 100 μm. **(C)** Quantification of experiment in **panel B**. Data represent the mCherry/GFP fluorescence intensity ratio averaged across all cells in a FOV, normalized by subtracting the background mCherry channel value. There was a higher mCherry/GFP fluorescence ratio in the light + Ca^2+^ compared to light – Ca^2+^ condition when 256 and 416 pulses were delivered (*N* = 6 FOVs per condition, 1-way ANOVA F(_4_,_25_)=25.43, *P* = 1.70e-8; Sidak’s multiple comparison’s test, \*\*\**P* < 0.001). **(D)** Relationship between duration of recording and FLiCRE activation. Fluorescence images of TRE-mCherry activation and GFP-CaM-uTEVp expression following light and E-stim (32 pulses at 20Hz every 3s) delivered for varying times. Scale bars, 100 μm. **(E)** Quantification of experiment in **panel D**. There was a higher mCherry/GFP fluorescence ratio in the light + E-stim condition compared to light – E-stim condition across all durations tested *(N* = 9 FOVs per condition, 2-way ANOVA interaction F_(4,64)_=10.63, *P* < 0.0001; Sidak’s multiple comparison’s test, \*\*\**P* < 0.001). **(F)** Testing the temporal resolution of FLiCRE. Light and E-stim were delivered either simultaneously for 10 min or staggered by 1 min in either direction. Fluorescence images of TRE-mCherry activation and GFP-CaM-uTEVp expression marker. Scale bars, 100 μm. **(G)** Quantification of experiment in **panel F**. The mCherry/GFP fluorescence ratio was higher in the simultaneous light and E-stim condition compared to the temporally staggered conditions (*N* = 9 FOVs per condition, 1-way ANOVA F_(2,24)_=72.6, *P* = 6.30e-11; Tukey’s multiple comparison’s test, \*\*\**P* < 0.0001). **See also Figures S1,S2.**

Having established a clear relationship between the number of action potentials fired and FLiCRE reporter expression, we then tested the temporal resolution of FLiCRE by staggering the delivery of light and E-stim in cultured neurons by 1 min in either direction. We observed TRE-mCherry FLiCRE labeling only when light and E-stim were simultaneously delivered, as opposed to offset by 1 min (**Figures 2F and 2G**). This was in contrast to our characterization of a similar tool, Cal-Light (Lee, 2017), which in our hands had lower signal-to-noise and lacked adequate temporal resolution to distinguish simultaneous versus sequential light and E-stim (**Figure S2**).

We then applied FLiCRE to the mammalian brain, one of the most challenging and complex of biological systems. To demonstrate that FLiCRE can detect elevated neural activity *in vivo*, we injected AAVs encoding FLiCRE and TRE-mCherry in the ventral tegmental area (VTA) of wildtype mice and implanted an optical fiber above the VTA to deliver blue light. To elevate calcium levels in the VTA, we injected mice with nicotine intraperitoneally (Wei et al., 2018) and delivered blue light to the VTA for 15 min. ~18 hrs later we analyzed the TRE-mCherry expression (**Figure 3A**). We observed more mCherry+ cells in mice treated with both nicotine and blue light, compared to just nicotine alone or just light alone (**Figures 3B and 3C**, and **Figures S3A and S3B**). These data demonstrate that FLiCRE can be used to record elevated neural activity *in vivo* during a single behavioral labeling session.

**Figure 3.**
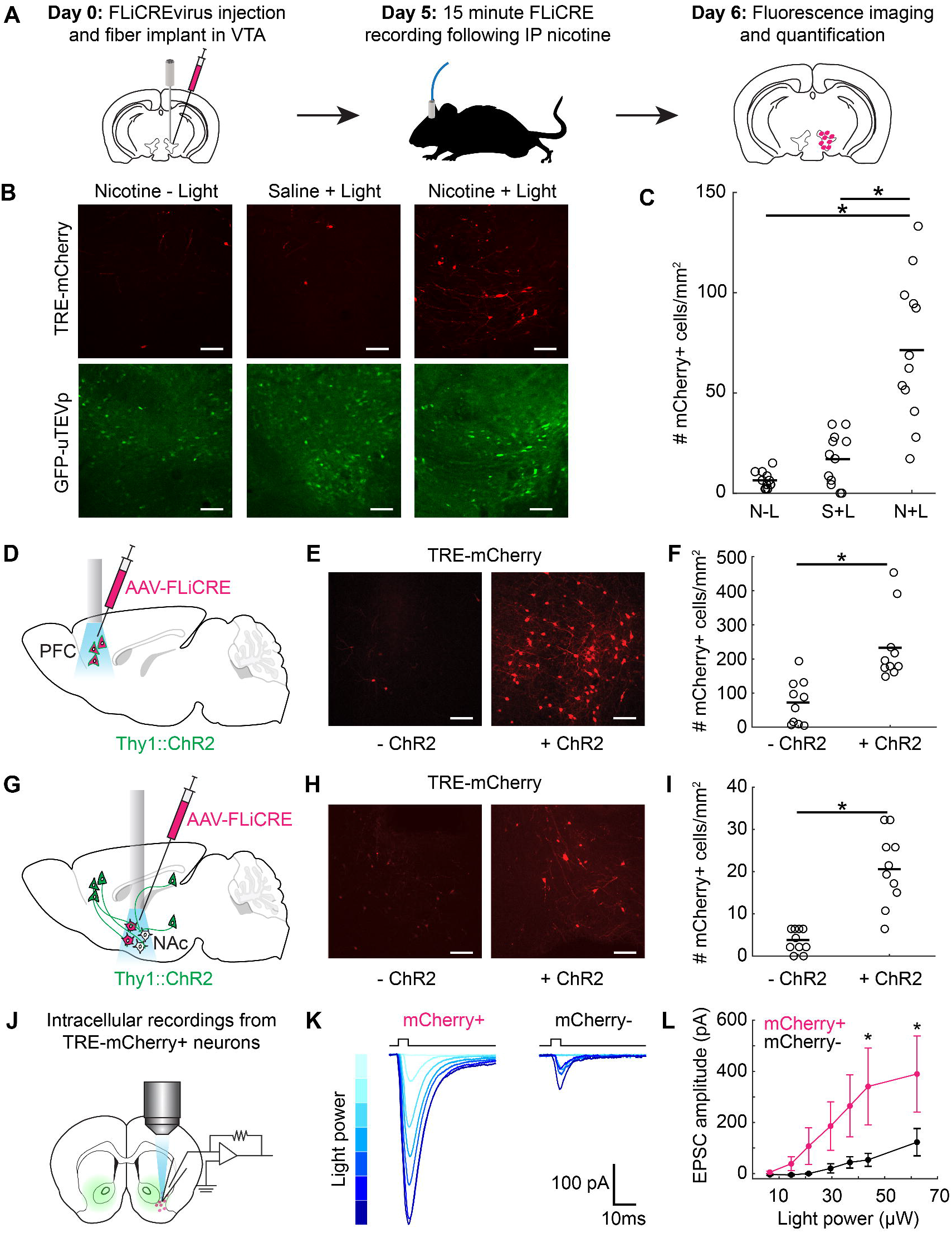
Application of FLiCRE to detect activated neuronal ensembles *in vivo.* **(A)** Timeline for *in vivo* FLiCRE injection, recording, and read-out in mice. hLOV1 FLiCRE viruses were injected in the VTA and an optical fiber was implanted above VTA. On day 5, mice were injected with 1.5mg/kg nicotine or saline intraperitoneally (IP), and 470nm light was delivered through an optical fiber to the VTA for 15 min. On day 6, mice were sacrificed. **(B)** Fluorescence images of FLiCRE TRE-mCherry reporter expression ~18 hrs following treatment with nicotine and blue light, or in control conditions omitting blue light (left) or nicotine (middle). Scale bars, 100 μm. **(C)** Quantification of experiment in **panel B**. Data show the mean number of mCherry+ cells/mm^2^ per FOV. There were more mCherry+ VTA cells in mice treated with nicotine and blue light compared to in control mice (*N* = 12 brain slices from 3 mice, each condition; 1-way ANOVA F_(2,33)_=, *P* =3.84e-8; Tukey’s multiple comparison’s test, \*\*\**P* < 0.0001). **(D)** Injection and optical fiber implant schematic for FLiCRE recording during optogenetic stimulation of PFC cell bodies. hLOV 1 FLiCRE viruses were injected in the PFC of Thy1::ChR2 transgenic mice, and an optical fiber was implanted in PFC. **(E)** Fluorescence images of TRE-mCherry FLiCRE reporter expression following 10 min of blue light delivered in ChR2+ and ChR2-(wildtype control) mice. Scale bars, 100 μm. **(F)** Quantification of experiment in **panel E**. Data show the mean number of mCherry+ cells/mm^2^ per FOV. There were more mCherry+ PFC cells in ChR2+ versus in ChR2-mice (*N* = 10 brain slices from 5 mice, each condition; Wilcoxon’s ranksum test, \**P* = 7.7e-4, U=150). **(G)** Injection and optical fiber implant schematic for FLiCRE recording of trans-synaptically activated neurons during optogenetic axonal stimulation. hLOV1 FLiCRE viruses were injected in the NAc of Thy1::ChR2 transgenic mice, and an optical fiber was implanted in NAc. **(H)** Fluorescence images of FLiCRE TRE-mCherry reporter expression following 10 min of simultaneous upstream axonal optogenetic stimulation and downstream somatic FLiCRE recording with blue light in ChR2+ and ChR2-mice. Scale bars, 100 μm. **(I)** Quantification of experiment in **panel H**. There were more mCherry+ NAc cells detected in ChR2+ versus in ChR2-mice *(N* = 10 brain slices from 5 mice, each condition; Wilcoxon’s ranksum test, \**P* = 2.9e-4, U=153). **(J)** Schematic for testing specificity of FLiCRE labeling of activated neurons using intracellular patching in acute *ex vivo* slices. An hLOV1 FLiCRE experiment was performed as in **panel G**. ~18 hrs later, patch clamp recordings were made from mCherry+ and mCherry-NAc neurons during brief pulses of blue light to elicit ChR2 axonal stimulation. **(K)** Example EPSCs from mCherry+ and mCherry-neurons recorded across increasing blue light powers. Light powers correspond to x-axis data points in **panel L**. **(L)** Quantification of experiment in **panel K**. mCherry+ NAc neurons had larger EPSCs than mCherry-NAc neurons at the light powers indicated (*N* = 6 mCherry+ and 5 mCherry-neurons; 2-way ANOVA interaction F_(6,54)_=2.44, *P* = 3.7e-2; Fisher’s LSD, \**P* < 0.05). **See also Figure S3.**

To further characterize the activity-dependence of FLiCRE while probing its labeling kinetics, we injected AAVs encoding FLiCRE and TRE-mCherry in the prefrontal cortex (PFC) of transgenic mice expressing the blue-light activated cation channel channelrhodopsin (ChR2 (Boyden et al., 2005)) chiefly in layer 5 excitatory neurons (Thy1::Chr2 (Arenkiel et al., 2007), **Figure 3D**). We implanted an optical fiber above the PFC to deliver blue light *in vivo*. The blue light simultaneously activates individual PFC neurons expressing ChR2 and uncages FLiCRE to enable recording and labeling of activated cells. We delivered 20Hz blue light pulses to the PFC for 10 min, and ~18 hrs later imaged PFC to detect TRE-mCherry expression. We observed more mCherry-expressing neurons in Thy1::ChR2 mice compared to control wildtype mice lacking the ChR2 gene (**Figures 3E and 3F**). We could also detect mCherry+ neurons activated during only 1 min of blue light stimulation, demonstrating the rapid labeling kinetics of FLiCRE (**Figures S3C–S3H**).

In the prior experiment, whenever blue light is delivered to stimulate ChR2 and elevate neural activity, FLiCRE is also activated. Thus to de-couple FLiCRE activation from ChR2 stimulation, we performed a variant of this experiment *in vivo* using a red-shifted excitatory opsin, bReaChES (Kim et al., 2016), instead of ChR2, to stimulate PFC neurons. This allowed us to confirm that the neuronal activity induced by opsin stimulation is not sufficient by itself to drive FLiCRE reporter expression in the absence of blue light. Even though this experiment required the injection of an additional virus encoding bReaChES, which potentially lowered the infection efficiency of all four components *in vivo*, we nevertheless observed lower FLiCRE reporter expression when we delivered only orange light to activate bReaChES, compared to when we delivered both blue light for FLiCRE and orange light for bReaChES (**Figures S3I–S3N**).

Given FLiCRE’s sensitive detection of neuronal activity, we asked whether this technology could be used to identify a subset of neurons *trans-synaptically* activated by stimulation of specific upstream axons. Such capability would represent a type of functional anterograde mapping, wherein we identify downstream neurons (not necessarily monosynaptically connected) that fire action potentials in response to stimulation of a genetically-defined upstream input. We chose to record which downstream neurons in the ventrolateral nucleus accumbens (NAc) are activated in response to temporally precise stimulation of local excitatory projection axons. Previous studies have shown that different inputs to this region drive varying behavioral outputs in rodents – for example, dopaminergic inputs can drive appetitive behaviors (Nicola et al., 2005; Steinberg and Janak, 2013), while excitatory glutamatergic inputs to different subregions can either increase (Britt et al., 2012; Stuber et al., 2011) or decrease (Kim et al., 2017; Qi et al., 2016; Reed et al., 2018) appetitive behaviors. This suggests that distinct sub-types of NAc neurons are targeted by different types of inputs, but technologies have been lacking to probe these anterograde connections *in vivo*.

To label the downstream neurons in NAc activated by excitatory glutamatergic inputs, we injected FLiCRE and implanted an optical fiber in the NAc of the Thy1::ChR2 mice. In these mice, ChR2 is expressed in excitatory layer 5 neurons in the frontal cortex (and in the hippocampus and amygdala) that send axonal projections to the NAc, but not in the NAc cell bodies themselves (**Figure 3G**). Blue light stimulation (20Hz pulses for 10 min) was delivered to simultaneously activate the upstream ChR2-expressing axons *and* uncage FLiCRE in NAc, giving labeling of NAc cells trans-synaptically stimulated by these long-range excitatory inputs. We observed more TRE-mCherry-expressing NAc neurons in ChR2+ mice compared to wildtype mice (**Figures 3H and 3I**). To test the specificity of this anterograde activity labeling, in a separate cohort of mice, we prepared acute brain slices of NAc for intracellular patch clamp recording after a similar FLiCRE recording (**Figure 3J**). We recorded excitatory postsynaptic currents (EPSCs) in mCherry+ and mCherry-NAc neurons in response to brief pulses of blue light, elicited by the ChR2-expressing axons present in the brain slices. We found that mCherry+ neurons exhibited larger EPSCs compared to mCherry-neurons, revealing that the FLiCRE-labeled neurons were more strongly synaptically-connected to ChR2-expressing excitatory inputs (**Figures 3K and 3L**).

Given that FLiCRE drives the expression of an exogenous reporter transcript (mCherry) in activated neurons, we reasoned that we should be able to detect FLiCRE activation via singlecell RNA sequencing (scRNA-seq). This would allow us to not only detect whether a cell had been activated during the FLiCRE recording, but also gain access to the cell’s native transcriptome and consequently cell typology. To our knowledge, only one prior study has reported detection of virally-targeted transcripts via scRNA-seq in adult neurons (Hanchate et al., 2018). To test whether we could detect FLiCRE activation via scRNA-seq, we performed the same FLiCRE recording in Thy1::ChR2 mice to label glutamate-activated NAc neurons with TRE-mCherry, but this time harvested (Saunders et al., 2018) the NAc to generate a barcoded single-cell RNA library (**Figure 4A**). We sequenced ~10,000 cells pooled across 3 experimental replicates from 5 mice. Dimensionality reduction and unsupervised clustering identified the broad classes of cell-types expected from this brain region (**Figures 4B–4D**). FLiCRE transcripts were enriched in the neuronal cell-types, as expected from use of the synapsin promoter (**Figures 4E and 4F**). Re-clustering of the identified neurons reported several expected cell-types in the NAc (Gokce et al., 2016; Muñoz-Manchado et al., 2018), including 3 subtypes of dopamine type 1 receptor-expressing medium spiny neurons (D1 MSNs), dopamine type 2 receptor medium spiny neurons (D2 MSNs), 2 interneuron subtypes, cholinergic subtypes, and 2 small additional clusters corresponding to cortical neurons included in the dissection that we omitted from further analysis (**Figures 5A and 5B**). These clusters were similarly represented across the three experimental replicates (**Figures S4A and S4B**), and they were also detected in samples from wildtype uninjected mice, importantly showing that the expression of FLiCRE AAVs did not grossly alter the transcriptome-based classification of cell-types in the brain (**Figures S4C and S4D**).

**Figure 4.**
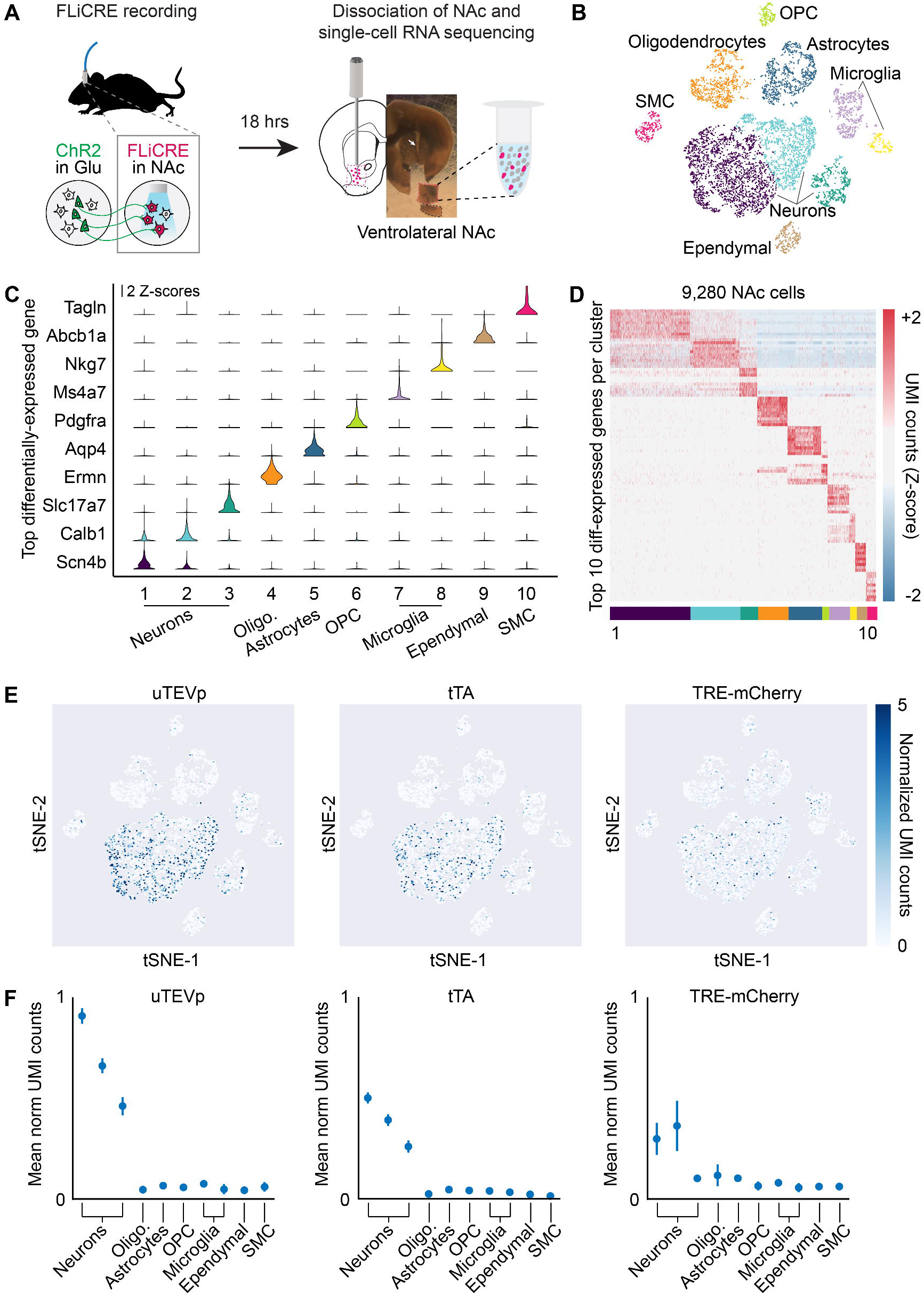
Detection of FLiCRE components using single-cell RNA sequencing. **(A)** Schematic for scRNA-seq following hLOV1 FLiCRE recording during afferent excitatory axon stimulation in NAc (as in **Figure 3E**). Single-cell libraries were generated using 10X Genomics kits and sequenced using Illumina Nextseq. **(B)** *t*-distributed stochastic neighbor embedding (t-SNE) of ~10,000 cells passing quality control metrics, colored by clustering analysis. Data clustered using unsupervised K-means clustering (cells pooled across 3 experimental replicates from 5 mice). **(C)** Violin plots of the top-differentially expressed gene identified per cluster. Y-axis represents Z-scored normalized UMI counts (see **STAR Methods**). **(D)** Heatmap of the top 10-differentially expressed genes identified per cluster. Normalized UMI counts for each gene were Z-scored across clusters. **(E)** Heatmap of normalized UMI counts detecting *uTEVp, tTA,* and *mCherry* across all cells plotted in t-SNE space. **(F)** Mean normalized UMI counts calculated for cells in each cluster type. FLiCRE transcripts were detected in cell clusters corresponding to neurons.

Among the NAc neuronal clusters, we asked whether the TRE-mCherry expression was enriched in any subset. We classified neurons expressing either *mCherry, tTA* (the transcription factor component of FLiCRE), or *uTEVp* (the protease component of FLiCRE) as having normalized UMI counts greater than 1 (**Figures 5C, S5A, and S5B**). As the expression of *tTA* and *uTEVp* varied across the different clusters, we calculated an enrichment score of *mCherry+* neurons (those active during the recording) relative to *uTEVp/tTA+* neurons (those infected with FLiCRE) – thus taking into account the differences in FLiCRE expression among the different clusters shown in **Figure 5C**. We found that *mCherry+* neurons were enriched in the D1 MSN1 sub-type compared to the other neuronal clusters (**Figure 5D**). This D1 MSN1 cluster was enriched in markers such as *Pdyn, Tac1, Wfs1, Rgs4,* and *Calb1* (**Figure 5E**). To further control for the differences in *uTEVp*/*tTA* expression across the neuronal clusters, we re-analyzed our dataset using only neurons that robustly expressed either the *uTEVp* or *tTA* FLiCRE components in their transcriptomes (**Figure 5F**). Re-clustering of these FLiCRE+ neurons revealed 3 different clusters, F1-F3. The *mCherry* reporter was enriched in only a single cluster, F1; the control components of *uTEVp* and *tTA* were similarly expressed across the three clusters, confirming that differences in FLiCRE expression were not driving the changes in mCherry enrichment across clusters (**Figure 5G**). We asked if cluster F1 represented a subset of any of the 7 broader neuronal clusters, and found that a majority of F1 neurons belonged to the D1 MSN1 cluster (**Figure 5H**), while clusters F2 and F3 likely represent a subset of the D1 MSN2 and D2 MSN clusters, respectively (**Figures S5C and S5D**). The *mCherry+* F1 cluster was also enriched with the same *Tac1* and *Calb1* marker genes as the D1 MSN1 cluster (**Figure 5I**). In addition, *mCherry+* and *FLiCRE+* neurons selected from our entire pool of neurons were also enriched with the *Tac1* and *Calb1* marker genes (**Figure S5E**), further confirming that FLiCRE activation primarily labeled this single NAc D1 MSN1 sub-type.

**Figure 5.**
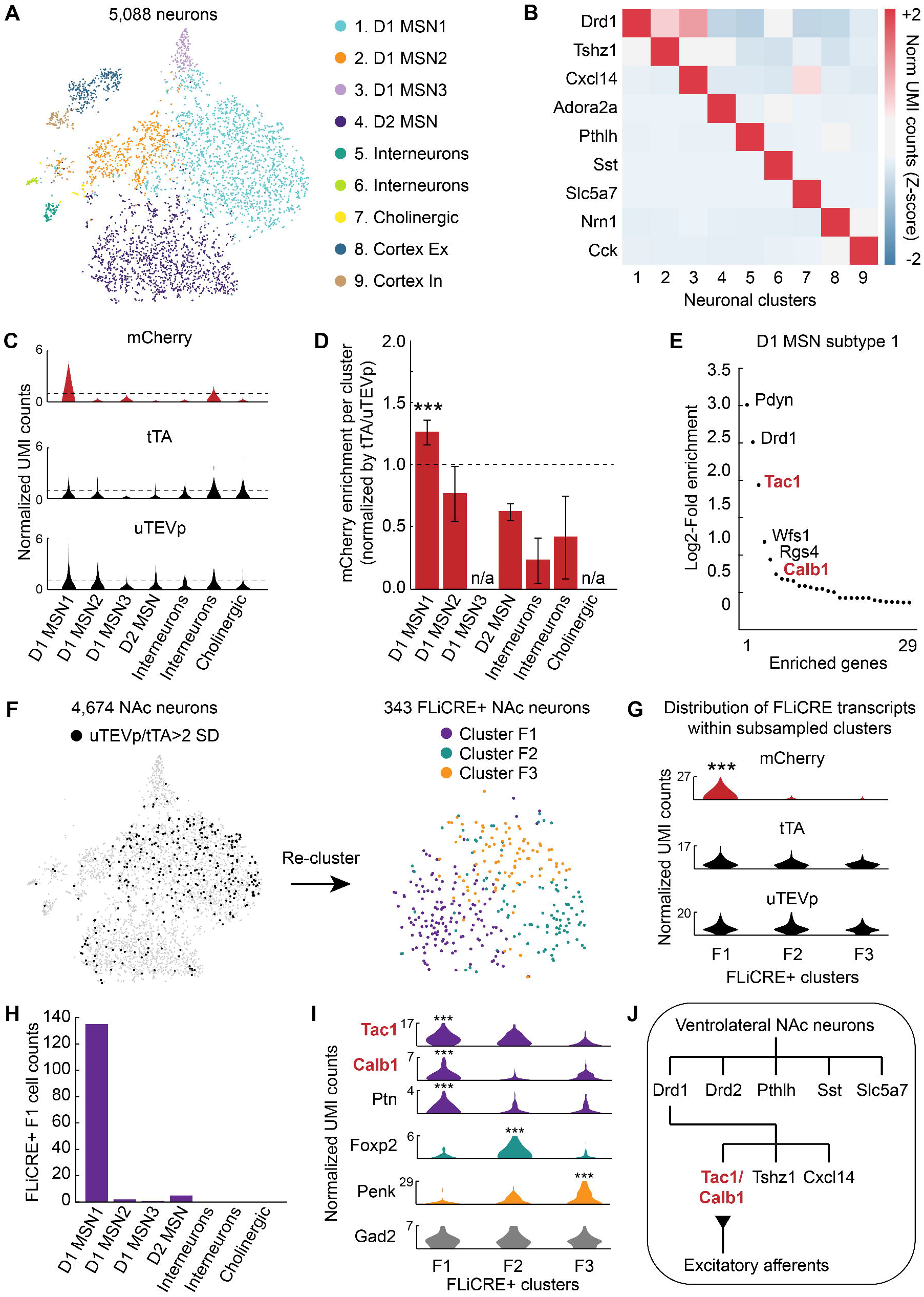
Simultaneous transcriptomic and calcium history read-out with FLiCRE. **(A)** t-SNE embedding of ~5,000 neurons, sub-clustered from cell clusters 1-3 in **Figure 4C**. Data re-colored and annotated across 9 identified neuronal clusters (K-means). **(B)** Heatmap of cell-type specific marker genes identified per cluster. Normalized UMI counts for each gene were Z-scored across clusters. **(C)** Violin plots of the distribution of normalized UMI counts of *mCherry, tTA,* and *uTEVp* within each cluster. The horizontal dashed line indicates a normalized UMI count of 1. **(D)** Enrichment of mCherry+ neurons, normalized by the number of FLiCRE+ neurons in each cluster (see **STAR Methods**). The D1 MSN1 cluster is enriched in mCherry+ neurons (*N* = 3 experimental replicates, Binomial test, 0.05 FDR corrected \*\*\**P* = 3.17e-8). **(E)** Enriched genes in the D1 MSN1 cluster compared to all other clusters. Y-axis represents the Log2 ratio of expression in neuronal cluster 1 compared to in all other cells. (Negative binomial test, corrected for multiple comparison’s using the Benjamini-Hochberg procedure, 0.1 FDR). **(F)** Left: t-SNE embedding of neurons from **panel A**, thresholded by *uTEVp* or *tTA* transcript expression. FLiCRE+ black cells have *uTEVp/tTA* UMI counts > 2 standard deviations (SD) of the mean. Right: t-SNE embedding of re-clustered FLiCRE+ neurons reveals 3 sub-types, F1-F3. **(G)** Violin plots of the distribution of normalized UMI counts of *mCherry, tTA,* and *uTEVp* within each sub-sampled cluster (Binomial test, 0.10 FDR corrected \*\*\**P* < 0.1). Y-axis represents normalized UMI counts. **(H)** The number of subsampled F1 cluster cells that belong to each broader neuronal cluster found in **panel A**. The majority of F1 cluster cells belong to D1 MSN subtype 1. **(I)** Violin plots of the distribution of all differentially expressed genes identified per subsampled cluster. Cluster F1 was enriched with *Tac1, Calb1,* and *Ptn;* Cluster F2 was enriched with *Foxp2;* and cluster F3 was enriched with *Penk* (Binomial test, 0.10 FDR corrected \*\*\**P* < 0.1). Violin plots for control gene *Gad2* are shown in grey. Y-axis represents normalized UMI counts. **(J)** Schematic of the summary of findings from the RNA-seq and FLiCRE experiment. **See also Figures S4 and S5.**

These sequencing findings suggest that incoming excitatory axons in the Thy1::ChR2 mice preferentially activate a distinct sub-type of downstream NAc neurons (**Figure 5J**). We next asked whether these downstream neurons might have a specific functional role in driving behavior related to motivation. We took advantage of the modular design of FLiCRE to express a red-shifted optogenetic cation channel bReaChES, instead of a static mCherry fluorophore, in activated NAc neurons; this configuration would allow us to subsequently re-activate these neurons *in vivo* using orange light. As before, we injected FLiCRE viruses into NAc of Thy1::ChR2 mice and performed the same 10-min stimulation and recording protocol, but now using a TRE-mCherry-p2a-bReaChES virus as the FLiCRE activation reporter (**Figure 6A**). To test whether these cells could play a functional role in driving behavior, we activated these neurons with orange light ~18 hrs after FLiCRE recording during a real-time place preference (RTPP) assay (**Figure 6B**).

**Figure 6.**
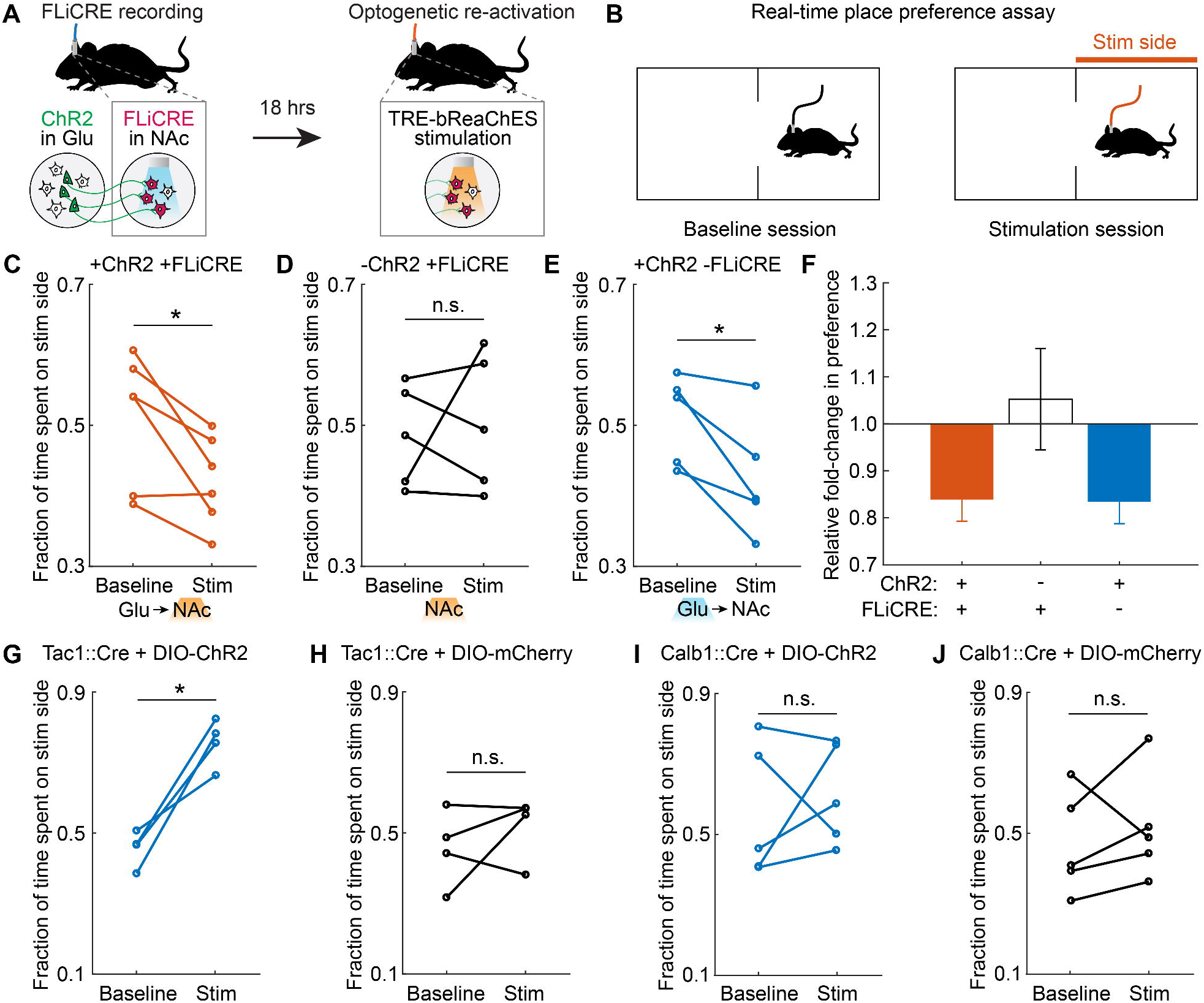
Control of behavioral function by manipulating previously-activated neuronal ensembles labeled with FLiCRE. **(A)** Schematic of experimental timeline. hLOV1 FLiCRE recording was performed during 10 min of excitatory axonal stimulation in NAc (as in **Figure 3E** using Thy1::Chr2 mice); but using aTRE-mCherry-p2a-bReaChES red-shifted excitatory opsin as the reporter. ~18 hrs later, orange light was delivered to re-activate the NAc neurons labeled during the FLiCRE recording. **(B)** Schematic of real-time place preference (RTPP) test. During the baseline session (Baseline), mice freely explore a two-sided chamber. During the stimulation session (Stim), light is delivered through the optical fiber when the mouse is on one side of the chamber (Stim side). The time spent on the stimulation side was compared during the Stim versus Baseline session. **(C)** RTPP results during orange light stimulation of TRE-bReaChES-expressing NAc cell bodies (labeled by FLiCRE during axonal stimulation of Thy1::Chr2 mice as in **panel A**). Mice avoided the Stim side (*N* = 6 mice; Student’s paired t-test, \**P* = 2.2e-2, t_(5)_=3.26). **(D)** RTPP results during orange light delivery to NAc cell bodies in wildtype mice (FLiCRE labels spontaneously active neurons). Mice do not consistently prefer or avoid the Stim side (*N* = 5 mice; Student’s paired t-test, n.s. *P* = 0.71, t_(4)_=0.41). **(E)** RTPP results during blue light stimulation of ChR2 excitatory afferent axons in NAc of Thy1::ChR2 mice (not expressing FLiCRE). Mice avoided the Stim side (*N* = 5 mice; Student’s paired t-test, \**P* = 2.7e-2, t_(4)_=3.41). **(F)** Summary of fold-change in preference for stimulation side from data in **panels C-E**. **(G,H)** Tac1::Cre mice were injected in NAc with either Cre-dependent ChR2 (DIO-ChR2) or DIO-mCherry, and then underwent the RTPP test during blue light delivery in NAc. ChR2-expressing mice preferred the Stim side (*N*_chR2_ = 4 mice, Student’s paired t-test, \**P* = 0.01, t_(3)_ = 5.71). Control mCherry-expressing mice exhibited no change in preference for the Stim side (*N*_mCherry_ = 4 mice, Student’s paired t-test, n.s. *P* = 0.41, t_(3)_ = 0.95). **(I,J)** Calb1::Cre mice were injected in NAc with either DIO-ChR2 or DIO-mCherry, and then underwent the RTPP test during blue light delivery in NAc. There was no change in preference for the Stim side in either cohort (*N*_ChR2_ = 5 mice, Student’s paired t-test, n.s. *P* = 0.61, t_(5)_ = 0.55; *N*_mCherry_ = 5 mice, Student’s paired t-test, n.s. *P* = 0.50, t_(3)_ = 0.74). **See also Figure S6.**

We found that orange light stimulation of the trans-synaptically activated NAc neurons, labeled by FLiCRE, drove a real-time place avoidance behavior in mice (**Figure 6C**). In contrast, orange light delivered into NAc of wildtype control mice, where the FLiCRE labeling was performed in the absence of excitatory axon stimulation, did not drive a consistent change in behavior (**Figure 6D**). In a separate cohort of Thy1::ChR2 mice, direct blue light activation of the long-range excitatory axons (expressing ChR2) in NAc also drove a real-time place avoidance behavior (**Figure 6E**), suggesting that activation of either the upstream or downstream nodes of this circuit is aversive to mice (**Figures 6F and Figures S6A–S6E**; see **Methods** for note on which specific subregion of NAc was targeted). We also performed two additional behavioral control experiments, the first in which Thy1::ChR2 mice received no blue light in NAc (and thus no NAc activation and no FLiCRE recording), but orange light was still delivered during the RTPP assay (**Figure S6F**). In the second control, Thy1::ChR2 mice received no blue light in NAc, but received optogenetic stimulation of the upstream excitatory ChR2-expressing cell bodies in the PFC (and thus still received NAc activation but no FLiCRE recording), followed by orange light during the RTPP assay (**Figures S6G-H**). In both controls, we did not observe a consistent change in behavioral preference (**Figures S6I–S6L**).

The cell-type manipulated by FLiCRE in this experiment is positive for both *Tac1* and *Calb1* markers, as shown by the single-cell RNA sequencing data (**Figure 5**). FLiCRE provides specific genetic access to this sub-type, which is not currently accessible by other methods (existing mouse lines or cell-type specific promoters). We asked whether stimulating a broader, less specific, set of NAc neurons that encompass the *Tac1/Calb1+* cells identified by FLiCRE, would elicit the same behavioral avoidance response. To test this, we injected Cre-dependent ChR2 into the NAc of Tac1::Cre or Calb1::Cre mice and performed RTPP assays with blue light stimulation. Activation of all *Tac1+* neurons in NAc drove an appetitive instead of aversive place preference response (**Figures 6G, 6H, S6M, and S6N**), while activation of all *Calb1+* neurons in NAc had no effect on place preference (**Figures 6I, 6J, S6O, and S6P**). These results demonstrate that the putative *Tac1*/*Calb1*+ neurons specifically labeled and activated using FLiCRE have a distinct functional role in behavior compared to the general population of all *Tac1+* or all *Calb1+* neurons.

We have demonstrated here that FLiCRE has enhanced sensitivity and kinetics compared to prior technologies both *in vitro* and *in vivo* (**Table S3**). However, a central challenge to all activity-dependent labeling tools, including FLiCRE, is background labeling of basally active neurons *in vivo*. As a first step to meeting this challenge, we present a proof-of-principle FLiCRE variant that employs a luciferin- and calcium-dependent gating of the LOV domain using a blue-light emitting luciferase (Kim et al., 2019) (**Figure 7A**). We fused the luciferase NanoLuc between the Calmodulin and uTEVp domains of FLiCRE (**Figure 7B**). In the presence of high calcium and NanoLuc’s luciferin substrate furimazine, the Calmodulin-NanoLuc-uTEVp will be brought into proximity of the membrane-bound component of FLiCRE to allow NanoLuc-LOV BRET. The uTEVp can then access TEVcs and release the transcription factor to the nucleus, as in the original FLiCRE design. The key difference between this new luciferin-gated design and the light-gated version of FLiCRE is that there is now a double-dependency on calcium for transcription factor release: both the LOV domain uncaging *and* the protease cleavage require high calcium in order to occur (**Figures 7C and 7D**). We hypothesized that this would generate a tool that is better at distinguishing between basal and stimulus-induced neural activity. To test this, we performed a luciferin- or light-gated FLiCRE experiment in either cortical or hippocampal cultured neurons, which have different levels of basal activity. In cortical neurons, where the spontaneous basal activity is very low (**Figure 7E**), both versions of FLiCRE reported high light ± activity and furimazine ± activity signal-noise ratios (**Figures 7F and 7G**). However, in hippocampal neurons which exhibit high basal calcium activity (**Figure 7H**), the light-gated version of FLiCRE could not distinguish between the basal vs. high calcium environments; only the luciferin-gated version produced a detectable furimazine ± activity signal-noise ratio (**Figures 7I and 7J**). This new luciferin-gated design principle could potentially reduce non-specific labeling of basally active neurons *in vivo,* while also removing the requirement of optical fiber implant and light delivery into the brain.

**Figure 7.**
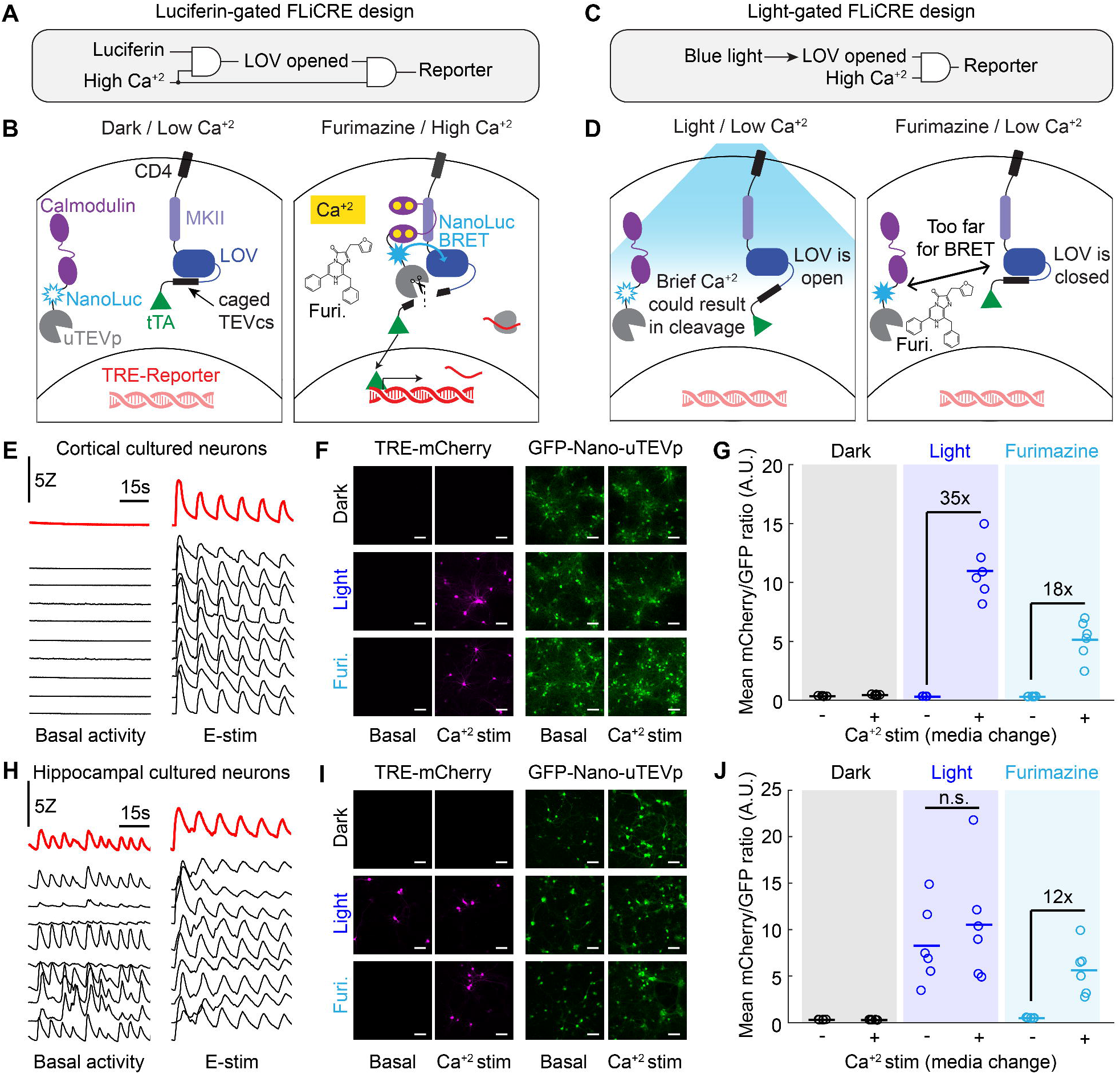
A versatile luciferin-gated FLiCRE design with steeper Ca^2+^ dependency. **(A)** Logic gate schematic of luciferase-gated FLiCRE design. LOV opening is dependent on both a luciferin and high Ca^2+^. **(B)** Mechanism of luciferase-gated FLiCRE. A blue luciferase, NanoLuc, is fused between CaM and uTEVp. When NanoLuc’s chemical substrate, furimazine, is supplied to the neurons during high Ca^2+^ activity, NanoLuc activates LOV via proximity-induced BRET. **(C)** Logic gate schematic of the blue light-gated FLiCRE design. LOV opening is dependent only on blue light and not on high Ca^2+^. **(D)** Schematic of the steeper Ca^2+^ dependency of luciferin-gated FLiCRE compared to light-gated FLiCRE. With blue light but low Ca^2+^ levels, the LOV domain is always open, and brief increases in intracellular Ca^2+^ (such as in unstimulated hippocampal neuron cultures) can cause uTEVp cleavage. However, with luciferin-gated FLiCRE, the LOV domain remains closed when there is low Ca^2+^ levels even when furimazine is present. **(E)** GCaMP transients of cortical neurons in untreated wells (left) and wells treated with E-stim (right; 50 pulses delivered at 20Hz every 10s). Black traces represent individual cells, red traces indicate entire FOV. **(F)** Fluorescence images of TRE-mCherry FLiCRE reporter and GFP-CaM-NanoLuc-uTEVp expression following 15 min of high Ca^2+^ stimulation (fresh media change) and blue light, or high Ca^2+^ and furimazine. Scale bars, 100 μm. **(G)** Quantification of experiment in **panel F**. Data represent the average mCherry to GFP fluorescence intensity ratio averaged across all cells in a FOV. There was a 35-fold increase in mCherry/GFP fluorescence ratio between light ± Ca^2+^ conditions (*N* = 6 FOVs; Wilcoxon’s signrank test, \**P* = 0.03, W=21). There was an 18-fold increase in mCherry/GFP fluorescence ratio between furimazine ± Ca^2+^ conditions (*N* = 6 FOVs; Wilcoxon’s signrank test, \**P* = 0.03, W=21). **(H)** GCaMP transients of individual hippocampal neurons in untreated wells (left) and wells treated with E-stim (right). Black traces represent individual cells, red traces indicate entire FOV. **(I)** Fluorescence images of TRE-mCherry FLiCRE reporter and GFP-CaM-NanoLuc-uTEVp expression following 15 min of high Ca^2+^ stimulation and blue light, or high Ca^2+^ and furimazine, as in **panel F**. Scale bars, 100 μm. **(J)** Quantification of experiment in **panel I**. There was not a significant fold difference in the mCherry/GFP fluorescence ratio between light ± Ca^2+^ conditions (*N* = 6 FOVs; Wilcoxon’s signrank test, *P* = 0.31, W=16). There was a 12-fold increase in mCherry/GFP fluorescence ratio between furimazine ± Ca^2+^ conditions (*N* = 6 FOVs; Wilcoxon’s signrank test, \**P* = 0.03, W=21).

## DISCUSSION

FLiCRE is a light-gated calcium integrator that provides stable genetic access to transiently activated cells. In contrast to real-time indicators of intracellular calcium (Chen et al., 2013; Tian et al., 2009), FLiCRE and other calcium integrators enable secondary interrogation of the molecular profiles of activated cellular ensembles as well as functional manipulation. FLiCRE is distinct from other calcium integrators in its speed and specificity (e.g., compared to FLARE (Wang et al., 2017) and Cal-Light (Lee, 2017), see **Figure S2**) and ability to functionally manipulate marked cells (e.g., compared to CaMPARI (Fosque et al., 2015)). Some studies have indirectly performed calcium integration by using the signal from real-time calcium indicators to select individual cells for photoconversion *ex vivo* (Lee et al., 2019) or opsin activation *in vivo* (Carrillo-Reid et al., 2019; Jennings et al., 2019; Packer et al., 2015; Rickgauer et al., 2014; Szabo et al., 2014), but these approaches are limited to smaller fields of view that must be targeted using sophisticated optical methodologies.

While calcium signaling is ubiquitous across biology, we chose here to apply FLiCRE to study the mammalian brain – specifically to investigate the molecular identity of an acutely activated ensemble of neurons, and to causally manipulate this subpopulation of neurons to influence behavior. Previous efforts have linked transcriptomics with neural activity by detecting elevated IEG expression in the endogenous transcriptome (Hrvatin et al., 2018; Moffitt et al., 2018; Wang et al., 2018), or using IEG promoter-driven transcriptional reporters to manipulate neurons previously activated during a stimulus (Allen et al., 2017; Guenthner et al., 2013; Liu et al., 2012). However, IEG expression is a less direct read-out of neuronal activity and has much slower dynamics compared to intracellular calcium (Hrvatin et al., 2018). Furthermore, IEG-based transcriptional tools rely on slow-metabolizing drugs to gate the time-window of recording, resulting in temporal resolution on the order of 4-6 hrs (DeNardo et al., 2019), in contrast to the 1 min temporal resolution of FLiCRE. Nevertheless, future studies directly comparing FLiCRE labeling with IEG labeling may be useful for understanding how IEG activity is linked to calcium activity.

Taking advantage of the specificity and sensitivity of FLiCRE, we identified a D1 MSN sub-type in the NAc that expresses *Tac1*, *Calb1*, and *Ptn*, and is activated *in vivo* by upstream excitatory axon stimulation. *Tac1* is considered a marker gene for all D1 MSNs (Lobo et al., 2006), *Calb1* is reported as a general MSN matrix marker gene (Liu and Graybiel, 1992), and *Ptn* has been reported to have drug-dependent regulation in the striatum in response to drugs such as levodopa or amphetamine (Ferrario et al., 2004; González-Castillo et al., 2015). This particular *Tac1/Calb1/Ptn+* D1 MSN subtype has not been explicitly described before in prior single-cell RNA sequencing datasets (Gokce et al., 2016; Stanley et al., 2019; Zeisel et al., 2018), although new cell-types co-expressing *Tac1* and *Penk,* a D2 MSN marker (Stanley et al., 2019) or *Drd2* and *Calb1* (Gokce et al., 2016) have been reported.

We found that FLiCRE-driven optogenetic stimulation of these putative *Tac1/Calb1/Ptn+* neurons elicited avoidance behavior in mice. This effect was specific to this sub-type, as optogenetic stimulation of either Tac1::Cre or Calb1::Cre NAc neurons did not drive avoidance behaviors in mice (currently there is no genetic Cre line for Ptn). While a more closely-matched control would have been to stimulate a similar number of *Tac1+* and *Calb1+* NAc neurons as those labeled by FLiCRE, this control is currently not feasible given the available transgenic lines. Furthermore, in this study we relied on the transgenic Thy1::ChR2 strain to label excitatory glutamatergic projections to NAc, which are reported to come from the prefrontal cortex, amygdala, and hippocampus. Future studies may identify which of these specific upstream excitatory projections is responsible for activating the FLiCRE-targeted *Tac1/Calb1/Ptn+* neurons we identified here; although prior studies have found that glutamatergic inputs to the NAc from all three areas appear to drive the same behaviors (Britt et al., 2012; Reed et al., 2018). As FLiCRE captures calcium spiking which can be elicited monosynaptically or polysynaptically, future studies will also be needed to elucidate the exact monosynaptic partners of this D1 MSN subtype.

Our behavioral findings, along with our sequencing results highlighting the heterogeneous transcriptional profile among different D1 MSN subtypes, suggest that finer granularity is required to define the functional neuronal subtypes in NAc. Indeed, prior studies using Cre lines to target *Drd1* or *Drd2* MSNs in the striatum have reported conflicting findings regarding causal roles in driving motivated behaviors. While it is thought that D1 MSNs drive appetitive behaviors and D2 MSNs drive avoidance behaviors (Kravitz et al., 2012; Lobo et al., 2010), recent evidence shows that both subtypes can drive either behavior, particularly in the NAc (Al-Hasani et al., 2015; Soares-Cunha et al., 2016; Soares-Cunha et al., 2019). These discordant reports may target different D1 or D2 MSN subtypes that are challenging to delineate using only genetic-targeting techniques, underscoring the importance of technologies like FLiCRE to access functionally-defined neuronal subpopulations.

We therefore envision that FLiCRE may enable a new class of biological inquiry into the transcriptomic and functional properties of acutely activated cell states. Future studies can couple FLiCRE transgene expression with *in situ* RNA sequencing technologies (Moffitt et al., 2018; Wang et al., 2018) to discover the spatial organization of functionally-defined ensembles. Using FLiCRE to drive the expression of a proximity labeling enzyme such as TurboID (Branon et al., 2018) could also enable activity-defined proteomic analysis downstream of the transcriptome. Additionally, driving the expression of actuators other than opsins, such as base editors (Anzalone et al., 2020) or synaptic silencers (Stachniak et al., 2014), may reveal other properties of functional circuits. The protease-based circuit that is the central feature of FLiCRE could potentially be adapted for other synthetic protease-based circuit designs (Fink et al., 2019; Gao et al., 2018).

We also demonstrated a new design principle for FLiCRE that relies on calcium-induced BRET between a blue light emitting luciferase NanoLuc and the LOV domain. This luciferin-gated version of FLiCRE not only exhibits a steeper calcium dependency compared to the light-gated version of the tool, but it would also only require a non-invasive injection of NanoLuc’s substrate furimazine rather than exogenously delivered blue light through an optical fiber. Promisingly, recent studies have demonstrated the *in vivo* use of NanoLuc in mice (Oh et al., 2019; Su et al., 2020) and in the brain (Germain-Genevois et al., 2016) to perform bioluminescence imaging. Further engineering and optimization of this luciferin-gated FLiCRE design will be needed to enable non-invasive, molecular recording of integrated calcium activity across the brain.

## ACKNOWLEDGEMENTS

We thank W. Allen and E. Richman (Stanford University) for assistance with initial scRNA-seq experiments; C. Ramakrishnan (Stanford University) and L. Ning (Stanford University) for assistance with cultured neurons; and J. Huguenard (Stanford University) for assistance with electrophysiology recordings during electric field stimulation. This research was supported by the Walter V. and Idun Berry Postdoctoral Fellowship Program (C.K.K.); EMBO long-term postdoctoral fellowship (ALTF 1022-2015; M.I.S.); NIMH (F32 MH115668; P.H.); NIMH and Stanford Psychiatry (L.E.F.); the Wu Tsai Neurosciences Institute and NIDA (P50 DA042012; R.C.M.); NIMH, NIDA, the Defense Advanced Research Projects Agency Neuro-FAST program, the NOMIS Foundation, the Wiegers Family Fund, the Nancy and James Grosfeld Foundation, the H. L. Snyder Medical Foundation, the Samuel and Betsy Reeves Fund, the Gatsby Foundation, the AE Foundation, and the Fresenius Foundation (K.D.); the Chan Zuckerberg Biohub and NIMH (R01 MH119353; A.Y.T.).

## AUTHOR CONTRIBUTIONS

C.K.K. performed *in vivo* experiments, single-cell RNA sequencing, and data analysis. M.I.S. performed protein engineering, cell culture experiments, and AAV1/2 virus production. P.H. performed intracellular patching experiments following FLiCRE labeling with guidance from R.C.M. L.E.F. performed intracellular patching experiments during electric field stimulation. A.Y.T. and K.D. supervised all aspects of the project. C.K.K., M.I.S., K.D., and A.Y.T. conceptualized the project, interpreted the data, and wrote the manuscript with edits from all authors.

## SUPPLEMENTAL FIGURE LEGENDS

**Figure S1.**
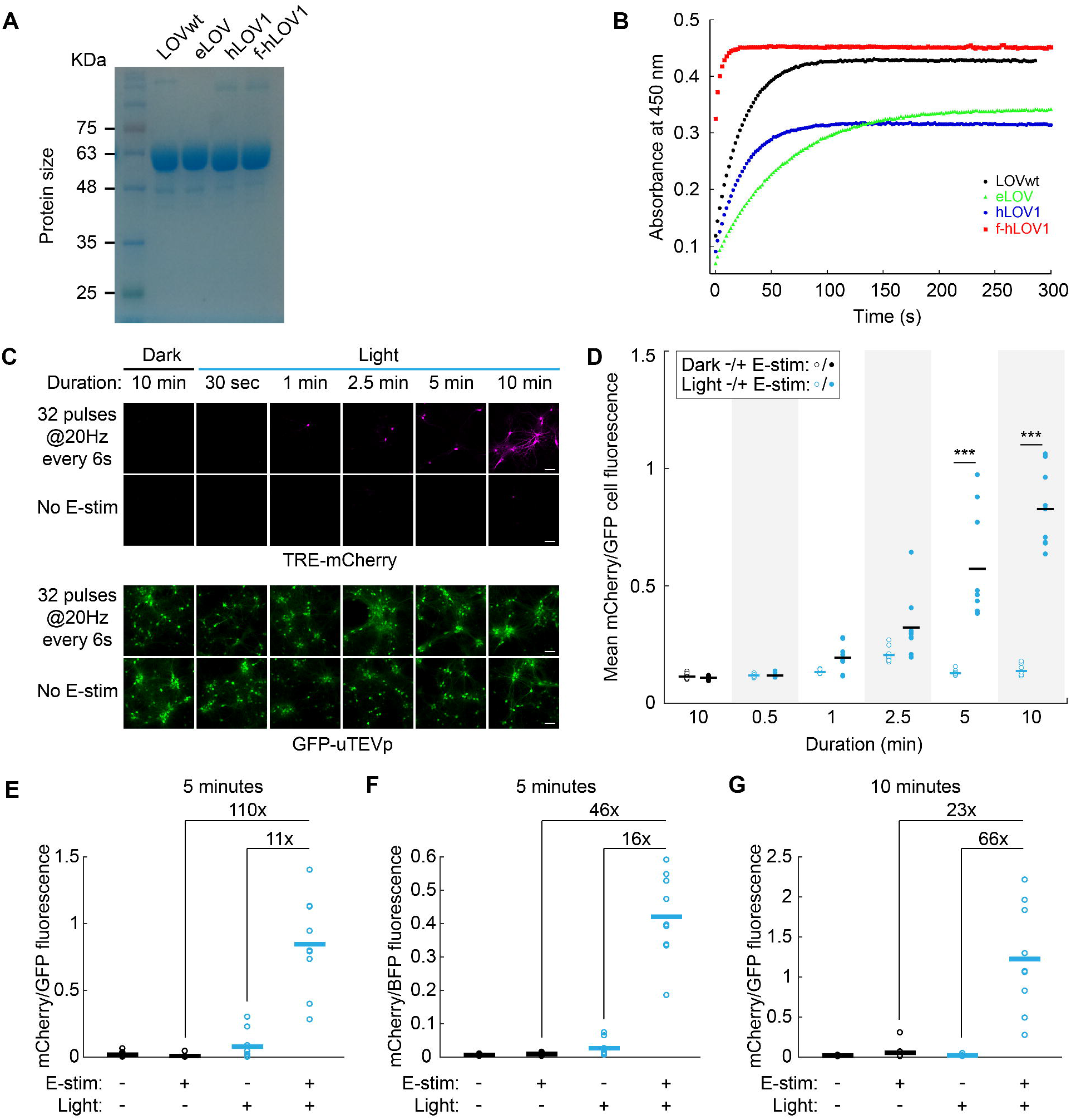
Related to Figures 1 and 2. Kinetics of different LOV domains and performance of FLiCRE in cultured neurons. **(A)** SDS-PAGE (9%) gel electrophoresis of purified LOV proteins stained by Coomassie Blue. **(B)** Recovery kinetics from the adduct state for the different LOV variants. **(C)** Repeat of experiment in **Figure 2D**, except the field stimulation is delivered less frequently (every 6s as opposed to every 3s). Fluorescence images of TRE-mCherry activation and GFP-calmodulin-uTEVp expression were taken ~18 hrs following light and electric field stimulation for the indicated times. Scale bars, 100 μm. **(D)** Quantification of experiment shown in **panel C**. There was a higher mCherry/GFP cell fluorescence ratio in the light + E-stim condition compared to light – E-stim condition during the 5 and 10 min recordings (*N* = 9 FOVs per condition, 2-way ANOVA interaction F_(4,64)_=33.62, *P* < 0.0001; Sidak’s multiple comparison’s test, \*\*\**P* < 0.001). **(E-G)** Cultured cortical neurons were infected with hLOV1 FLiCRE viruses in 3 different biological replicates as in **Figure 2**. Neurons were treated with blue light and electric field stimulation (32 pulses delivered at 20Hz every 3 s, 1-ms pulse width) for either 5 min or 10 min. Neurons in **panel F** were infected with BFP-Calmodulin-uTEVp, while neurons in **panels E** and **G** were infected with GFP-Calmodulin-uTEVp. Significant fold-change increases in signal are displayed above the data. The mCherry channel was normalized by subtracting the mean background value of each image outside of the cell masks. Light ± E-stim SNRs ranged from 11x to 66x, while E-stim ± Light SNRs ranged from 23x to 110x.

**Figure S2.**
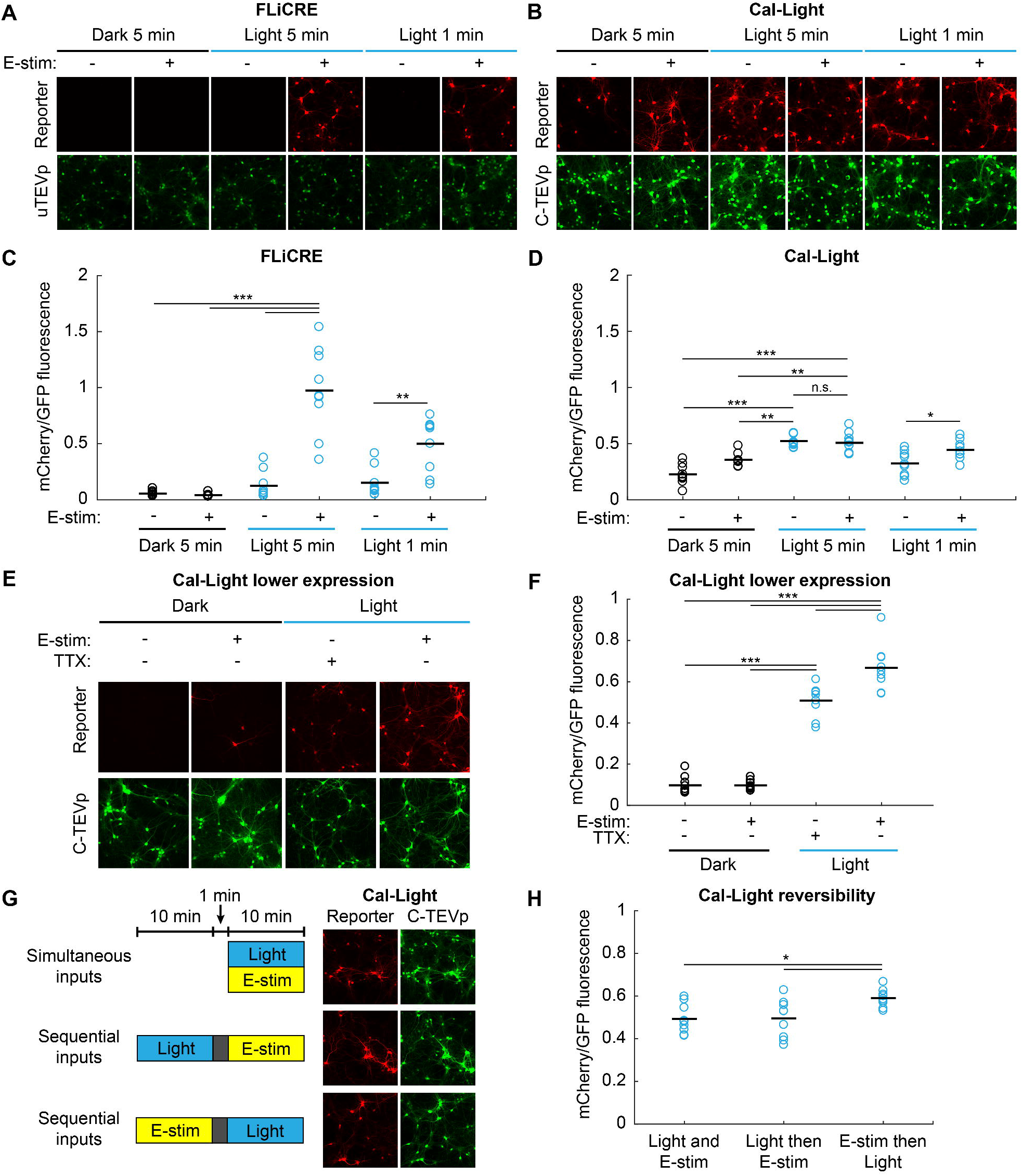
Related to Figure 2. Comparison of FLiCRE to Cal-Light in cultured neurons. **(A,B)** Side-by-side comparison of hLOV1 FLiCRE and Cal-Light (Lee, 2017) performance in cultured cortical neurons. Fluorescence images of TRE-mCherry activation and GFP-calmodulin-uTEVp (FLiCRE) or M13-C-TEVp-p2a-GFP (Cal-Light) expression were taken ~18 hrs following light and electric field stimulation (32 pulses delivered at 20Hz every 3 s, 1-ms pulse width). Scale bars, 100 μm. **(C,D)** Quantification of experiments shown in **panels A,B**. FLiCRE performance was similar to **Figure S1**, with a higher mCherry/GFP cell ratio in the “Light + E-stim” condition compared to the control conditions (*N* = 9 FOVs per condition, 2-way ANOVA interaction F_(2,48)_=21.54, *P* = 2.11e-7; Tukey’s multiple comparison’s test, \*\*\**P* < 0.001, \*\**P* < 0.01). For Cal-Light, we did not observe activity-dependence with the 5 min light +/− E-stim condition, and observed a small but significant fold change with 1 min of light +/− E-stim (*N* = 9 FOVs per condition, 2-way ANOVA interaction F_(2,48)_=4.42, *P* = 0.017; Tukey’s multiple comparison’s test, \*\*\**P* < 0.001, \*\**P* < 0.01, \**P* < 0.05). **(E)** Repeat of Cal-Light experiment at an earlier timepoint with lower expression of the tool (Day 4 following viral infection instead of Day 6). To match the parameters described in the paper (Lee, 2017), we used cultured hippocampal neurons and treated them with TTX in the “-E-stim” condition. Fluorescence images of TRE-mCherry activation and M13-C-TEVp-p2a-GFP expression were taken ~18 hrs following light and electric field stimulation (32 pulses delivered at 20Hz every 3 s, 1-ms pulse width). **(F)** Quantification of experiment shown in **panel E**. The “Light + E-stim” condition had a higher mCherry/GFP cell ratio than all other conditions; although the “Light – E-stim” still had a higher mCherry/GFP cell ratio compared to the Dark control conditions (*N* = 9 FOVs per condition, 2-way ANOVA interaction F_(1,32)_=11.16, *P* = 0.0021; Tukey’s multiple comparison’s test, \*\*\**P* < 0.001). **(G)** Testing the temporal resolution of Cal-Light in cultured cortical neurons. Light and E-stim were delivered either simultaneously for 10 min or staggered by 1 min in either direction (e.g. Light then E-stim, or E-stim then Light). Fluorescence images are shown for TRE-mCherry activation and M13-C-TEVp-p2a-GFP expression marker. Scale bars, 100 μm. **(H)** Quantification of experiment shown in **panel G**. The mCherry/GFP cell ratio was slightly higher in the sequential “E-stim then Light” condition compared to the other conditions (*N* = 9 FOVs per condition, 1-way ANOVA F_(2,24)_=5.75, *P* = 0.0091; Tukey’s multiple comparison’s test, \**P* < 0.05).

**Figure S3.**
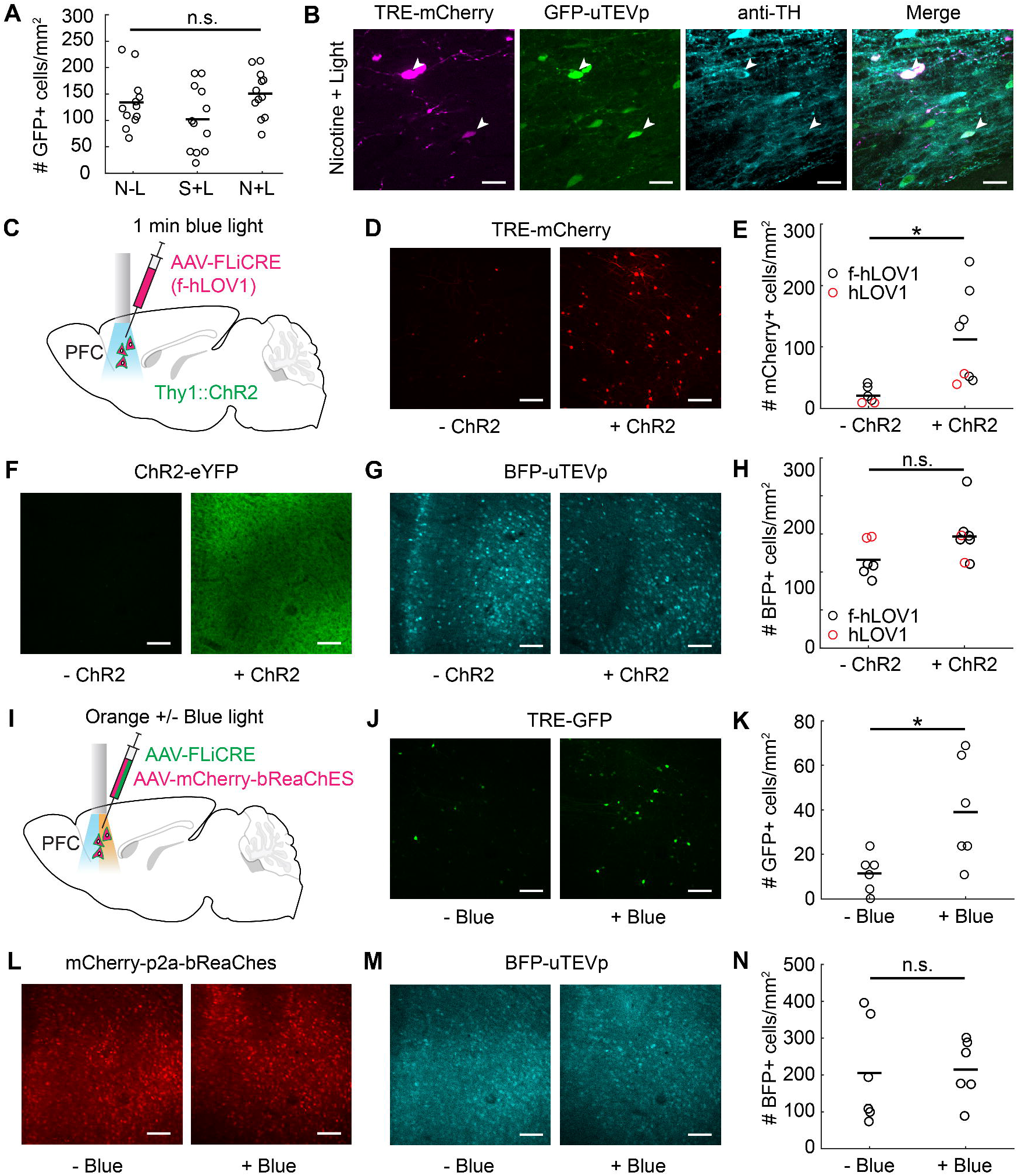
Related to Figure 3. Additional controls and quantification for *in vivo* FLiCRE experiments. **(A)** Quantification of the GFP-uTEVp expression shown in the nicotine FLiCRE recording experiment in **Figure 3B**. There was no difference in the number of GFP+ VTA cells detected across conditions (*N* = 12 brain slices from 3 mice, each condition; 1-way ANOVA F_(2,33)_ = 2.66 *P* = 0.085). **(B)** Fluorescence images of a 40x FOV in the VTA taken from a nicotine+light FLiCRE recording condition. Anti-TH staining was performed in a subset of slices to confirm targeting to the VTA. White arrows point to example TRE-mCherry FLiCRE reporter neurons that are positive for anti-TH staining for dopamine neurons. Further experiments must be performed to quantify which cell-types are labeled by FLiCRE recording in the VTA. Scale bars, 3 μM. **(C)** Injection and optical fiber implant schematic to enable FLiCRE recording during 1-min direct optogenetic stimulation of cell bodies. FLiCRE viruses (as in **Figure 3D**, except using f-hLOV1 instead of hLOV1) were injected in the PFC of Thy1::ChR2 transgenic mice, and an optical fiber was implanted in PFC. **(D)** Fluorescence images of FLiCRE TRE-mCherry reporter expression ~18 hrs following 1 min of simultaneous ChR2 optogenetic stimulation and FLiCRE recording. Wildtype control mice lacking ChR2 underwent the same surgical and experimental protocol. Scale bars, 100 μm. **(E)** Quantification of experiment shown in **panel D**. Data show the mean number of mCherry+ cells/mm^2^ per FOV. There were more mCherry+ PFC cells detected in ChR2+ mice compared to in wildtype control mice (*N*_+chR2_ = 8 brain slices from 4 Thy1::ChR2 mice, *N*_-chR2_ = 6 brain slices from 3 wildtype mice; Student’s unpaired t-test, \**P* = 0.014, t_(12)_=2.89). For each condition, 1 mouse was injected with FLiCRE hLOV1 instead of f-hLOV1 (red circles). **(F,G)** Fluorescence images of eYFP (**F**) and BFP-uTEVp expression **(G)** in wildtype control (-ChR2) and Thy1::ChR2 (+ChR2) mice shown in **panel D**. Scale bars, 100 μm. **(H)** Quantification of BFP-uTEVp expression shown in **panel G**. There was no difference in the number of BFP+ PFC cells detected in ChR2+ mice compared to in wildtype control mice (*N*_+ChR2_ = 8 brain slices from 4 Thy1::ChR2 mice, *N*_-ChR2_ = 6 brain slices from 3 wildtype mice; Student’s unpaired t-test, n.s. *P* = 0.083, t_(12)_=1.89). **(I)** Injection and optical fiber implant schematic to enable hLOV1 FLiCRE recording during 10-min direct optogenetic stimulation of cell bodies using a red-shifted excitatory opsin instead of ChR2. AAV-synapsin-mCherry-p2a-bReaChES and FLiCRE viruses (as in **Figure 3D**, except using TRE-GFP instead of TRE-mCherry) were injected in the PFC of wildtype mice, and an optical fiber was implanted in PFC. **(J)** Fluorescence images of FLiCRE TRE-GFP reporter expression ~18 hrs following 10 min of simultaneous orange light (for bReaChES optogenetic stimulation) and blue light (for FLiCRE recording; + Blue). Control mice underwent the same surgical protocol, but were exposed to 10 min of only orange light and not blue light (-Blue). Scale bars, 100 μm. **(K)** Quantification of experiment shown in **panel J**. There were more GFP+ PFC cells detected in mice given both orange and blue light (+Blue) compared to in control mice given only orange light (-Blue) (*N* = 6 brain slices from 3 mice, each condition; Student’s unpaired t-test, \**P* = 0.023, t(10)=2.68). **(L,M)** Fluorescence images of mCherry-p2a-bReaChES (**L**) and BFP-uTEVp expression **(M)** in control (-Blue) and experimental (+Blue) mice shown in **panel J**. Scale bars, 100 μm. **(N)** Quantification of BFP-uTEVp expression shown in **panel M**. There was no difference in the number of BFP+ PFC cells detected in the +Blue mice compared to in the -Blue control mice (*N* = 6 brain slices from 3 mice, each condition; Student’s unpaired t-test, n.s. *P* = 0.90, t_(10)_=0.13).

**Figure S4.**
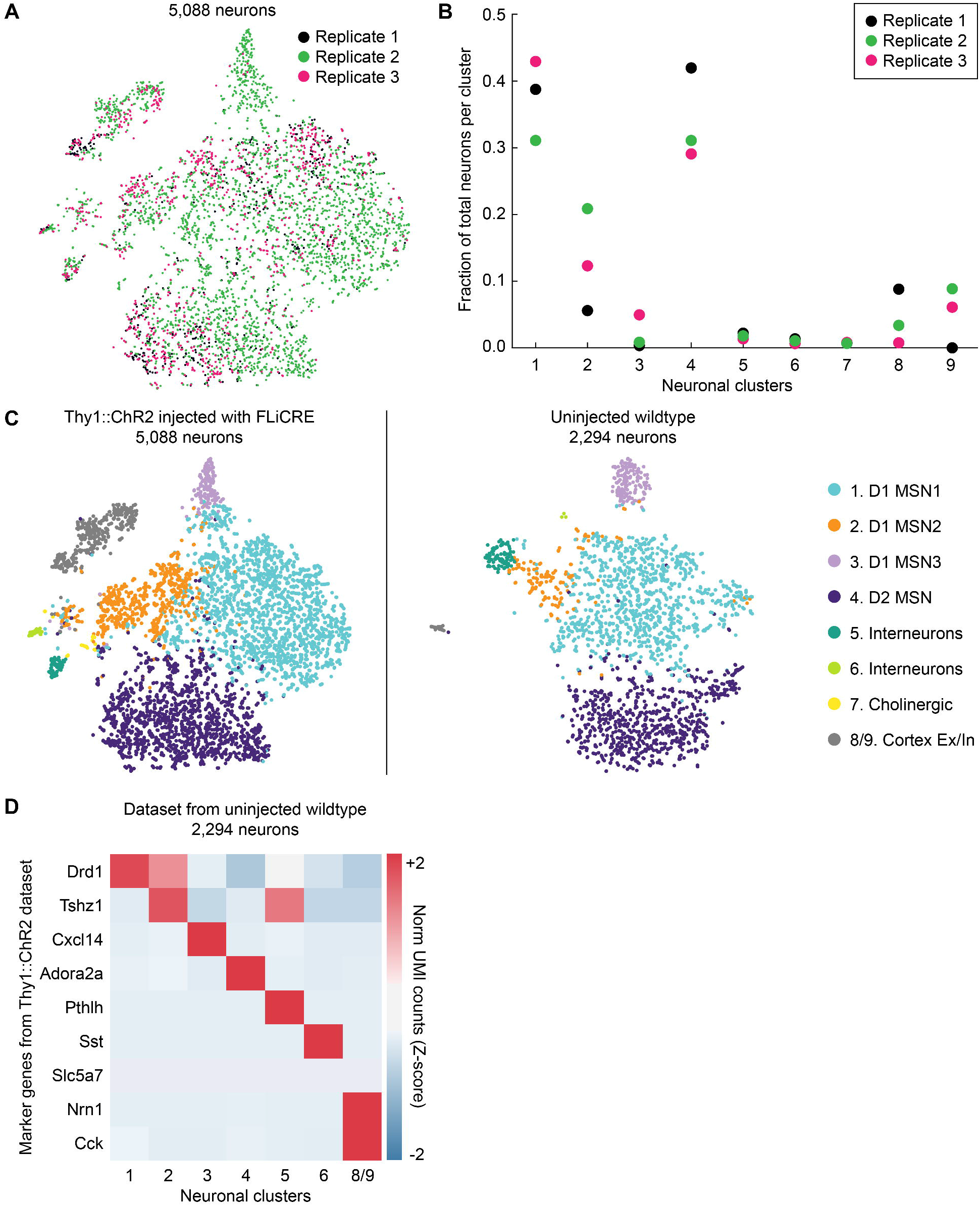
Related to Figure 5. Controls for scRNA-seq data. **(A)** t-SNE embedding of neurons from **Figure 5A**, color-coded by each of the 3 experimental replicates (pooled from 5 mice). **(B)** Fraction of total neurons in each cluster, calculated for each experimental replicate. Mice had similar distributions of neurons represented in each cluster. Clusters correspond to **Figure 5A**. **(C)** Side-by-side visualization of t-SNE embedding of the dataset analyzed in **Figure 5A** (Thy1::ChR2 mice injected with FLiCRE) and a separate dataset collected from an uninjected wildtype mouse (pooled from 2 technical replicates). **(D)** Heatmap of normalized UMI counts from the uninjected wildtype dataset, calculated for the top marker genes identified from the Thy1::ChR2 dataset in **Figure 5B**. The same neuronal clusters were identified across the two datasets, except that the rare *Slc5a7* cluster was not present in the uninjected wildtype dataset.

**Figure S5.**
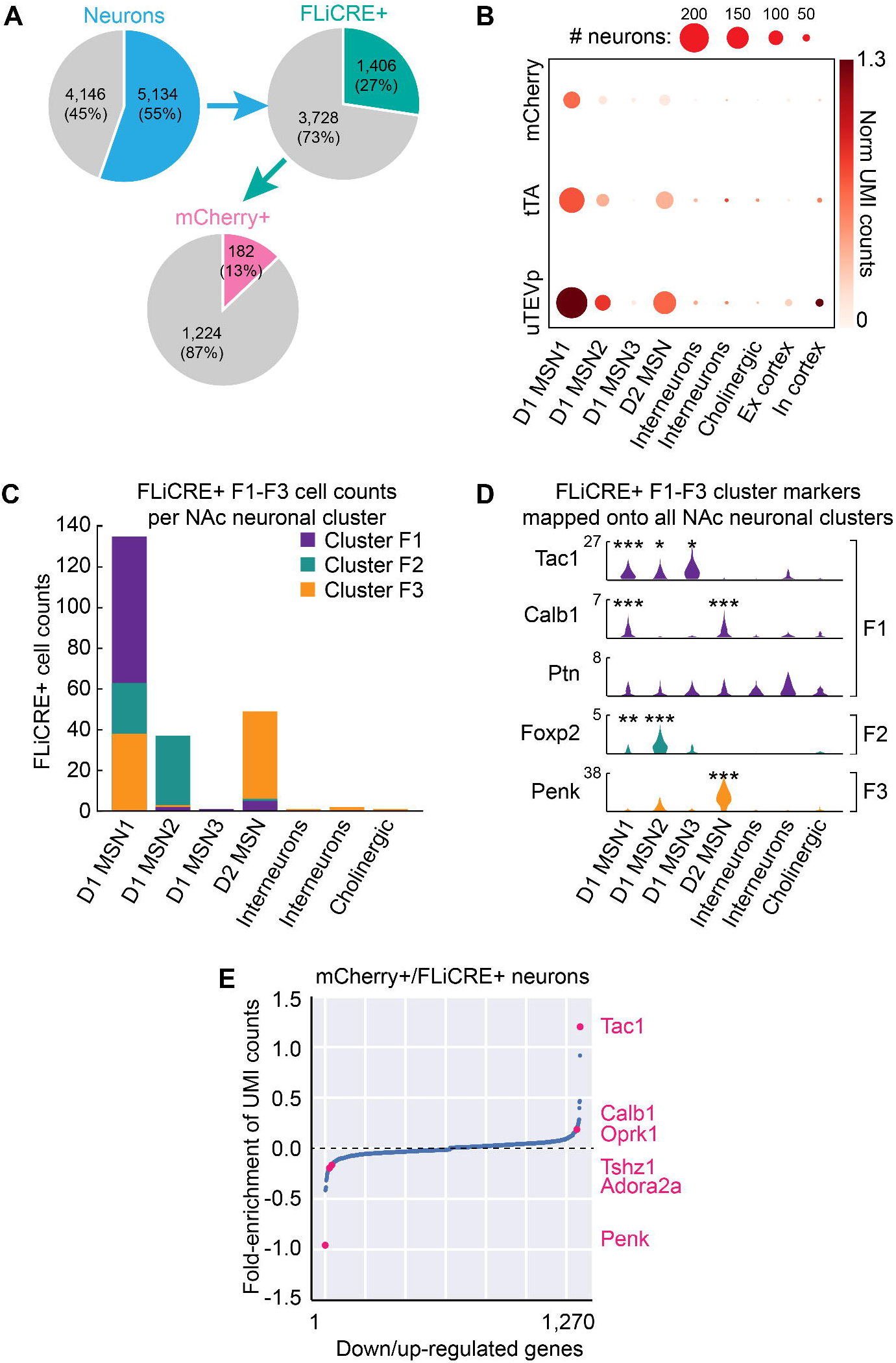
Related to Figure 5. Additional analysis of FLiCRE expression among scRNA-seq data. **(A)** Pie charts showing the percentages of all cells that are neurons, the percentages of neurons that express FLiCRE (> 1 normalized UMI count for either *uTEVp* or *tTA),* and the percentage of FLiCRE+ neurons that express mCherry (>1 normalized UMI count for *mCherry).* **(B)** Dotplot visualization of *uTEVp, tTA,* and *mCherry* transcript expression across all neurons averaged within each cluster. The size of the dots corresponds to the number of neurons with a normalized UMI count > 1, while the color corresponds to the mean normalized UMI count across all neurons in that cluster. Clusters 8 and 9, representing a small number of cortical neurons included in the dissection, were removed from further analysis in the paper. Across the 3 biological replicates, Cluster 8 was only present in 2 of the biological replicates, with an average of 3 mCherry+ neurons present in the cluster. Cluster 9 was present in all 3 biological replicates, with an average of 2 mCherry+ neurons present in the cluster. We assume these few mCherry+ neurons are due to background labeling. **(C)** Overlaid bar plots of FLiCRE+ F1-F3 cell counts per NAc neuronal clusters 1-7. F1-F3 clusters are those as in **Figure 5G**, and NAc neuronal clusters 1-7 are those as in **Figure 5A**. **(D)** Violin plot of the distribution of normalized UMI counts across neurons within each cluster. The marker genes represent those that are up-regulated in each of the FLiCRE+ NAc clusters identified in **Figure 5G**. Neuronal cluster 1 is enriched with *Calb1* and *Tac1*, and expression of *Ptn* is also detectable. Neuronal cluster 2 is enriched in *Foxp2*. Neuronal cluster 4 is enriched in *Penk (N* = 3 experimental replicates, Binomial test, 0.10 FDR corrected \*\*\**P* < 0.001, \*\**P* < 0.01, \**P* < 0.1). **(E)** Genes either up-regulated or down-regulated in mCherry+ and FLiCRE+ neurons (thresholded normalized UMI count>1), compared to 10,000 randomly shuffled distributions of cell labels. Both *Tac1* and *Calb1* were enriched in mCherry+/FLiCRE+ neurons.

**Figure S6.**
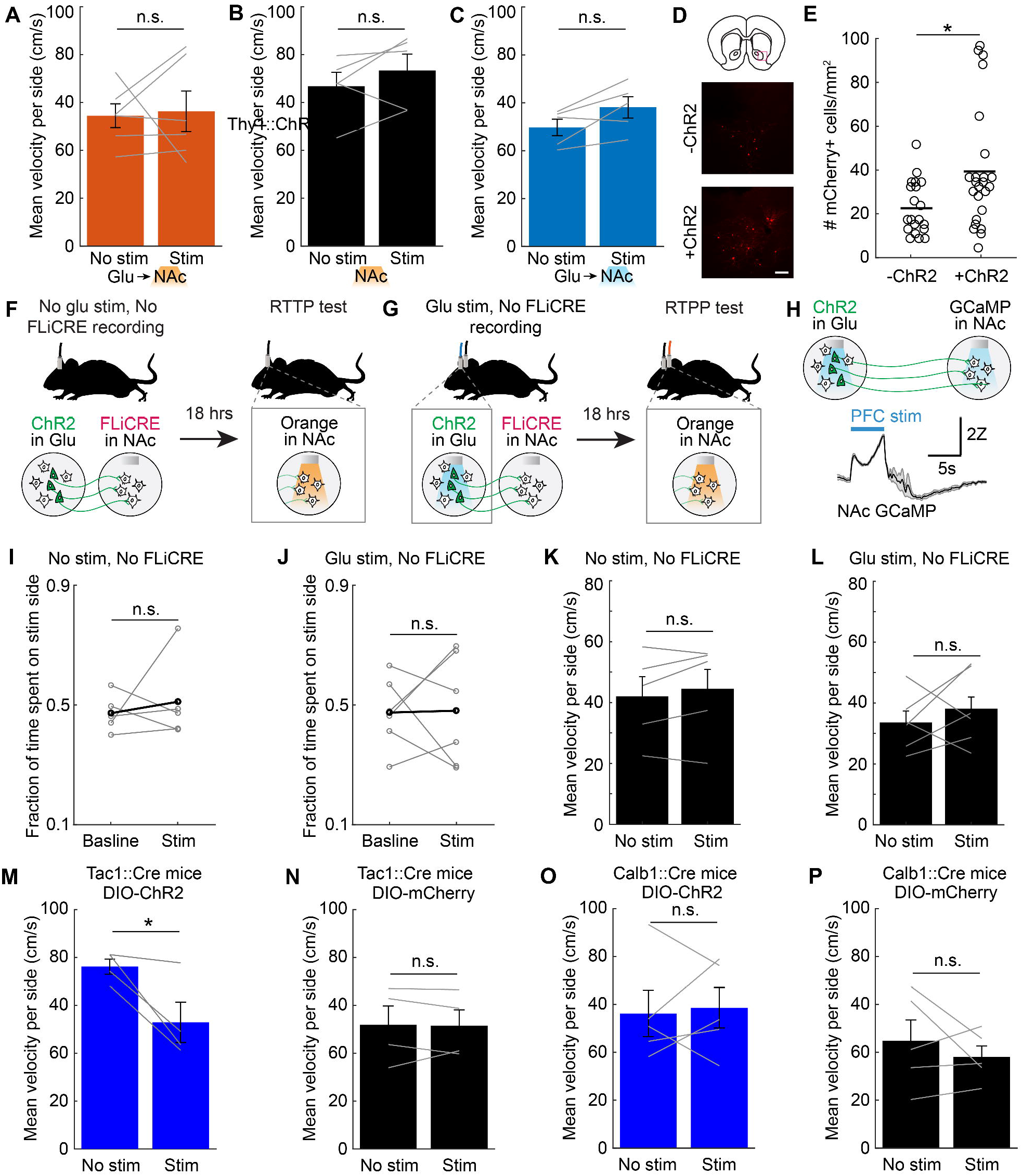
Related to Figure 6. Additional optogenetic behavior controls. **(A-C)** Velocity of mice measured during the Stim session of the real-time place preference assay from **Figures 6C**, **6D**, and **6E,** respectively. There was no significant difference in velocity on the non-stimulation side versus the stimulation side across experimental cohorts (*N*_(A)_ = 6 mice, Student’s paired t-test, n.s. *P* = 0.85, t_(5)_ = 0.20; *N*_(B)_ = 5 mice, Student’s paired t-test, n.s. *P* = 0.28, t_(4)_ = 1.26; *N*_(C)_ = 5 mice, Student’s paired t-test, n.s. *P* = 0.067, t_(4)_ = 2.50). **(D)** Fluorescence images of FLiCRE TRE-mCherry-p2a-bReaChES reporter expression in mice that underwent the real-time place preference test in **Figures 6C and 6D**. Scale bars, 100 μm. **(E)** Quantification of experiment shown in **panel D**. There were more mCherry-p2a-bReaChES+ NAc cells detected in ChR2+ mice compared to in wildtype ChR2-control mice (*N* = 24 brain slices from 6 ChR2+ mice and 20 brain slices from 5 wildtype mice; Wilcoxon’s ranksum test, \**P* = 3.1e-2, U=632). **(F)** Schematic describing a “-Activity/-Light” behavioral control for the real-time place preference test experiment in **Figure 6**. Thy1::ChR2 mice were injected with hLOV1 FLiCRE and implanted with an optical fiber in NAc as in **Figure 6A**. On day 5 following surgery, no blue light was delivered to the brain. On day 6, mice underwent a real-time place preference assay using orange light in NAc. **(G)** Schematic describing a “+Activity/-Light” behavioral control for the real-time place preference test experiment in **Figure 6**. Thy1::ChR2 mice were injected with hLOV1 FLiCRE in NAc as in **Figure 6A**, but an optical fiber was implanted above both the NAc and the PFC. Instead of delivering blue light to the NAc on day 5, blue light was delivered to the PFC to optogenetically stimulate ChR2-expressing cell bodies. This also causes downstream increased activity in NAc cell bodies, without initiating FLiCRE recording in NAc. The following day, mice underwent a real-time place preference assay using orange light in NAc. **(H)** *In vivo* GCaMP5G fiber photometry recording of NAc cell body bulk calcium responses to PFC optogenetic stimulation. ~5mW (measured at the optical fiber tip) was used for the PFC stimulation while only ~5μW was used for NAc GCaMP imaging (to minimize cross-stimulation of ChR2-expressing axons). Photometry trace represents the mean ± s.e.m. of five stimulation trials. Time-locked blue light stimulation NAc responses were replicated across 2 mice. **(I,J)** Results of real-time place preference assay for control experiments shown in **panels F** and **H.** There was no change in preference for the stimulation side during the Stim session compared to the Baseline session in either cohort (*N*_(I)_ = 5 mice, Student’s paired t-test, n.s. *P* = 0.52, t_(4)_ = 0.63; *N*_(J)_ = 6 mice, Student’s paired t-test, n.s. *P* = 0.94, t_(5)_ = 0.076). **(K,L)** Velocity of mice measured during the Stim session of the real-time place preference assay from **panels I** and **J**. There was no significant difference in velocity on the non-stimulation side versus the stimulation side in either cohort (*N*_(K)_ = 5 mice, Student’s paired t-test, n.s. *P* = 0.31, t_(4)_ = 1.17; *N*_(L)_ = 6 mice, Student’s paired t-test, n.s. *P* = 0.46, t_(5)_ = 0.81). **(M,N)** Velocity of Tac1::Cre mice measured during the Stim session of the real-time place preference assay from **Figures 6G and 6H**. There was a significant decrease in velocity on the stimulation side versus the non-stimulation side in ChR2-expressing Tac1::Cre mice **(M)**, but not in control mCherry-expressing Tac1::Cre mice **(N)** (*N*_ChR2_ = 4 mice, Student’s paired t-test, \**P* = 0.048, t(3) = 3.25; *N*_mCherry_ = 4 mice, Student’s paired t-test, n.s. *P* = 0.90, t_(3)_ = 0.13). **(O,P)** Velocity of Calb1::Cre mice measured during the Stim session of the real-time place preference assay from **Figures 6I and 6J**. There was no significant difference in velocity on the non-stimulation side versus the stimulation side in either cohort (*N*_ChR2_ = 5 mice, Student’s paired t-test, n.s. *P* = 0.79, t_(4)_ = 0.28; *N*_mCherry_ = 5 mice, Student’s paired t-test, n.s. *P* = 0.43, t_(3)_ = 0.89).

## STAR METHODS

### RESOURCE AVAILABILITY

#### Lead Contact

Further information and requests for resources and reagents should be directed to and will be fulfilled by the Lead Contact, Alice Ting.

#### Materials Availability

Plasmids generated in this study have been deposited to Addgene (**Table S1**). Single-cell RNA sequencing data generated in this study have been deposited to NCBI GEO (GSE160486).

#### Data and Code Availability

The datasets/code supporting the current study are available at www.github.com/kimck/FLiCRE-analysis-scripts and are available from the corresponding author on request.

### EXPERIMENTAL MODEL AND SUBJECT DETAILS

#### HEK293T cells

HEK293T cells (ATCC) were cultured as a monolayer in complete Dulbecco’s Modified Eagle Medium (DMEM, Corning), supplemented with 10% Fetal Bovine Serum (FBS, VWR) and 1% (v/v) penicillin-streptomycin (Corning, 5000 units/mL penicillin and 5000 μg/mL streptomycin) at 37°C, 5% CO_2_. This cell line has not been authenticated.

#### Cultured rat neurons

All protocols were approved by Stanford University’s Institutional Animal Care and Use Committee. Cortical neurons were harvested from male and female rat embryos at post embryonic day 18 following euthanasia by CO_2_ asphyxiation, as previously described (Loh et al., 2016), and described again below. Dissected cortical tissue was digested in papain (Worthington) and DNase I (Roche) for 30 min at 37°C. The cell suspension was filtered through a 40 μm nylon cell strainer. Cells were cultured in 24-well plates at 37°C, 5% CO_2_ either on a glass coverslip (0.1 mm thick) or directly on the plate. Neurons were grown in complete neurobasal media (Gibco) supplemented with 2% (v/v) B27 (Life Technologies, 1% (v/v) Glutamax (Life Technologies), 1% penicillin-streptomycin (VWR, 5 units/mL penicillin, 5 ug/mL streptomycin). At DIV4, 300 μL media was removed from each well and replaced with 500 μL complete neurobasal media supplemented with 10 μM 5-fluorodexoyuridine (FUDR, Sigma-Aldrich) to inhibit glial cell growth. Subsequently, approximately 30% of the media in each well was replaced with fresh complete neurobasal media every 3 days.

#### Mouse animal models

All experimental and surgical protocols were approved by Stanford University’s Institutional Animal Care and Use Committee. For FLiCRE experiments, 7-8 week old male/female wildtype C57BL/6J (Jackson Laboratory Strain 000664) and homozygous Thy1::ChR2-YFP (Jackson Laboratory Strain 007612) transgenic mice were used. For real-time place preference control experiments, 7-8week old male/female Tac1-IRES2-Cre (Jackson Laboratory Strain 021877) and Calb1-IRES2-Cre (Jackson Laboratory Strain 028532) mice were used. Mice were group-housed and given *ad libitum* food and water prior to surgery. Mice were maintained on a reverse 12-hour light/dark cycle for 1 week prior to behavioral assays.

### METHOD DETAILS

#### Plasmid constructs and cloning

Constructs were cloned into the pAAV viral vector (a gift from F. Zhang, MIT). **Table S1** lists all plasmids used in this study. For all constructs, PCR fragments were amplified using Q5 polymerase (NEB), and vectors were double-digested with NEB restriction enzymes and ligated to gel-purified PCR products using Gibson assembly. Ligated plasmids were introduced into competent XL1-Blue bacteria via heat shock transformation.

#### HEK293T FLiCRE experiments

For immunofluorescence imaging assays, cells were plated in 48-well plates pre-treated with 50μg/mL human fibronectin (Millipore). DNA transfections were performed with polyethyleneimine max according to the manufacturer’s specifications (PEI, Polysciences 24765). At ~80% confluency, we delivered the following DNA mixture into each well: 20 ng UAS-mCherry, 30 ng Calmodulin-TEVp, 50 ng CD4-MKII-LOV-TEVcs-Gal4 component, and 0.8 μl PEI max. Cells were incubated wrapped in foil overnight for 8-9 hrs.

The following day, cells were treated with blue light and calcium. For dark conditions, cells were handled under red light, which does not activate the LOV domain. Intracellular calcium was elevated using a final concentration of 2 μM ionomycin and 6 mM CaCl_2_. Blue light was delivered using a custom-built 467-nm LED light box, that delivered 60 mW/cm^2^ light at a 33% duty cycle (2s light every 6 s). After incubation for the indicated time, the solution was removed in each well and replaced with fresh 200 μl complete DMEM. Cells were kept wrapped in foil in the incubator for an additional 8-9 hrs and then imaged under the confocal.

#### Purification of LOV variants and kinetics measurements

In a pYFJ16 vector, the *Avena sativa* phototropin1 LOV2 (404–560) domain was cloned as a fusion with a GS linker to the C-terminus of the *E. coli* maltose binding protein (MBP). Mutations were made by standard molecular biology techniques. All proteins were expressed in *E. coli* BL21 (DE3) cells grown in LB media with Ampicillin and Chloramphenicol at 37°C to an OD_600_ of 0.6 and induced with 1 mM IPTG. Cultures were then incubated for 18 hrs at 18°C. The fusion protein was purified by metal affinity chromatography (IMAC). Bacteria were spun down at 6,000 rpm for 6 min. The pellet was lysed with B-PER (20 mL/culture liter), mixed gently by inversion and spun down at 10,000 rpm for 15 min; the supernatant was incubated with Ni-NTA agarose beads in binding buffer (Tris 50 mM, NaCl 300 mM, pH 7.8) for 10 min. The slurry was placed in a peptide column and washed with washing buffer (Imidazole 30 mM, Tris 50 mM, NaCl 300 mM, pH 7.8). The protein was eluted with elution buffer (Imidazole 200 mM, Tris 50 mM, NaCl 300 mM, pH 7.8). The collected volume (intense bright yellow) was transferred to a centrifugal filter Amicon Ultra-15 and washed 3x with DPBS. The purity was analyzed by SDS–PAGE and Coomassie Blue staining. The purified protein was used right away.

#### UV-Vis spectroscopy of LOV proteins

We characterized the reversion kinetics of the LOV protein from light state to dark state following previous procedures (Zayner et al., 2012), described again here. UV-Vis spectra were acquired using a Nanodrop2000c spectrophotometer with a 5-nm bandwidth and a 1-cm pathlength cuvette at 22°C. Kinetic traces were acquired after photosaturation of freshly purified LOV protein with a MaestroGen UltraBright LED transilluminator (470 nm). Samples (~10 μM in DPBS) were irradiated for 30 seconds and the absorbance at the λ_max_ (~448 nm) was measured every 2 seconds.

#### Immunofluorescence imaging and analysis

Confocal imaging was performed with a Zeiss AxioObserver inverted microscope with a 10x air objective. The following combinations of laser excitation and emission filters were used for various fluorophores: eGFP (491 nm laser excitation; 528/38 nm emission), mCherry (561 nm laser excitation; 617/73 nm emission), BFP (405 nm excitation; 450/30 nm emission). Acquisition times ranged from 100 to 500 ms. All images were collected with SlideBook (Intelligent Imaging Innovations) and processed with Fiji (Schindelin et al., 2012). For each FOV collected per condition, individual cells were first identified in the GFP channel (uTEVp expression). The mean UAS-or TRE-mCherry reporter expression was then calculated for each cell identified, and the mean mCherry/GFP value was calculated for each cell in the FOV. The mCherry/GFP cell ratios were then averaged across each FOV collected. Cell identification and quantification was performed using the CellSegm toolbox (Hodneland et al., 2013) in MATLAB R2017a (Mathworks).

#### AAV1/2 virus production, concentration, and titering

AAV1/2 viruses were prepared and concentrated as previously reported (Konermann et al., 2013; Wang et al., 2017) and described again here. HEK293T cells were grown to ~90% confluency in 3-4 T150 flasks per construct. Each flask was transfected with the following: 5.2 ug vector of interest, 4.35 ug AAV1 plasmid, 4.35 ug AAV2 plasmid, 10.4 ug DF6 AAV helper plasmid, and 130 μl PEI. Cells were incubated for 48 hrs, and then the supernatant media (30 mL total) was harvested for infection of cultured neurons.

To purify and concentrate the virus, the remaining HEK293T cells were pelleted at 800g for 10 min, and then resuspended in 20 mL of 150 mM TBS (150 mM NaCl, 20 mM Tris, pH = 8.0). 10% sodium deoxycholate (in water, Sigma) was added to the resuspended cells at a final concentration of 0.5%, along with 50 units/mL of benzonase nuclease (Sigma). The cell suspension was incubated at 37°C for 1 hour, and then cleared by centrifugation at 3,000g for 15 min. The supernatant was loaded to a HiTrap heparin column (GE Healthcare), pre-equilibrated with 10 mL of TBS, using a peristaltic pump (Gibson MP4). Following loading of the virus, the column was washed with 20 mL of 100 mM TBS using the pump, followed by 1mL of 200 mM TBS and 1mL of 300 mM TBS using a syringe. The virus was eluted using 1.5 mL 400 mM TBS, 3.0 mL 450 mM TBS, and then 1.5 mL of 500 mM TBS. The virus was then concentrated using a 15mL centrifugal unit (100K molecular weight cut off, Amicon), spinning at 2,000g for ~1 min, to a final volume of 500uL. The virus was further concentrated in a 1.5 mL centrifugal unit (Amicon) to ~100 μl.

To titer the concentrated viruses, 2 μl of virus was incubated with 1 μl DNase I (NEB, 2 units/μl), 4uL DNAse I buffer, and 33 μl H_2_O for 30 min at 37°C. The DNaseI was inactivated at 75°C for 15 min. 5 μl of this reaction was then added to 14 μl H_2_O and 1uL of Proteinase K (Thermo Fisher, 20 mg/mL) for 30 min at 50°C. Proteinase K was inactivated at 98°C for 10 min. 2 μl of this reaction was used for the qPCR reaction, along with 5uL of SYBR Green master mix (2x), 0.06 μl each of the forward and reverse primers (0.3 μM), and 2.88 μl of H_2_O. Primers were designed against the synapsin promoter and WPRE:

Syn-forward: 5’-GGGTGCCTACCTGACGAC
Syn-reverse: 5 ‘-GTGCTGAAGCTGGCAGTG
WPRE-forward: 5 ‘-CTGTTGGGCACTGACAATTC
WPRE-reverse: 5 ‘-AGAATCCAGGTGGCAACATA

Standardized curves were generated using purified/linearized AAV DNA plasmids that contained synapsin and WPRE. Three different standard curves were generated, for 0.05 ng, 0.1 ng, or 0.2 ng of linearized DNA. The titer of each viral sample was calculated in reference to the standard curve as fully described in ref (Wang et al., 2017).

#### Rat neuron culture FLiCRE experiments

Cortical neurons were cultured as described above. At 4 days in vitro (DIV4), 10 μM 5-fluorodexoyuridine (FUDR, Sigma-Aldrich) was added to the media to inhibit glial growth. The media was changed every 3 days with fresh complete neurobasal media (without FUDR). For the experiment shown in **Figure 7F**, hippocampal neurons were used instead of cortical neurons.

At DIV11-12, neurons were infected with a mixture of crude supernatant AAV1/2 viruses (see **Table S2**). Plates were wrapped in aluminum foil and kept in the incubator for an additional 6 days. On FLiCRE recording days, blue light was delivered as described above for HEK293T cell experiments. To elevate intracellular calcium, neurons were treated with electric field stimulation using two platinum iridium alloy wires (70:30, alfa-Aesar), bent into two parallel rods that were placed above the neurons at the edge of the wells. Current was delivered through these electrodes using a stimulator isolate unit (Warner, SIU-102b). To control the pulses, a Master 8 (AMPI) was used. Generally, we delivered 32 pulses at 20 Hz every 3 s, with 1-ms pulse width and the stimulator set to 48 mA. Imaging was performed the following day.

#### Electrophysiology during electric field stimulation

Recordings of neurons prepared and transfected as above were obtained in room temperature Tyrode’s medium (in mM: 150 NaCl, 4 KCl, 2 MgCl_2_, 2 CaCl_2_, 10 d-glucose, 10 HEPES, adjusted to pH 7.3-7.4 with NaOH, 320-330 osmolarity) supplemented with Tetrodotoxin (TTX; Tocris, approximately 1 μM when bath applied) where indicated, with a standard internal solution (in mM: 130 K-gluconate, 10 KCl, 10 HEPES, 10 EGTA, 2 MgCl_2_, to pH 7.25 with KOH, 300-310 osmolarity) in 3-6 MΩ glass pipettes. All data collected from whole-cell recordings. Recordingswere made using a Axoclamp-2B amplifier (Molecular Devices). Field current stimulation was applied through two platinum wires shaped to the recording chamber and placed approximately 1 cm apart. 45 mA stimuli were generated using a Warner SIU-102

Bipolar Stimulator in bipolar mode and pulse width, timing, and frequency were controlled with the recording software. Data were filtered at 4 kHz and digitized at 10 kHz with a Digidata 1440A analog-digital interface (Molecular Devices). pClamp10.6 software (Molecular Devices) was used to record and analyze data.

#### Mouse stereotaxic surgeries

For stereotaxis surgeries, mice were anesthetized with 1.5%–2.0% isoflurane, and placed in a stereotaxic apparatus (Kopf Instruments) on a heating pad. The fur was cut from the scalp and a midline incision was made. 3% hydrogen peroxide was applied to the skull, and a craniotomy was made above the injection site. Virus was injected using a 33-gauge beveled needle and a 10 μL Hamilton syringe (World Precision Instruments), controlled by an injection pump (Harvard Apparatus). 1,000 nL of virus was injected at 150 nL min^-1^. For FLiCRE recordings, a 400-μm core diameter, 0.39-NA optical fiber (Thorlabs) was implanted above the VTA, mPFC, or NAc. Implants were secured to the skull using dental adhesive (Parkell, C&B metabond). The following coordinates (in mm relative to bregma) were used for viral injections: VTA −3.2 A/P, 0.35 M/L, −4.6 D/V; mPFC +1.98 A/P, 0.35 M/L, −2.25 D/V; NAc +1.15 A/P, 1.6 M/L, −4.6 D/V. Fiber implants were placed at the same A/P and M/L locations, but 0.2 mm more dorsally. The virus titers, dilutions, and volumes used for each experiment are listed in **Table S2**.

#### Subregion of NAc targeted for FLiCRE experiments

For our FLiCRE experiments, we targeted the ventrolateral NAc, rather than the more commonly targeted core subregion. Excitatory afferents to the core are well-known to drive strong preference and appetitive behaviors (Britt et al., 2012; Stuber et al., 2011; Tye, 2012). However, prior studies have also shown that excitatory afferents into the shell region can drive avoidance behaviors or reduce appetitive behaviors (Kim et al., 2017; Qi et al., 2016; Reed et al., 2018). Thus, our findings that excitatory afferents into the ventrolateral NAc do not conflict with the existing literature. In addition, in our own hands, stimulation of the excitatory afferents in the NAc core of the Thy1::ChR2 mice drive the expected place preference behavior (data not shown).

#### Mouse FLiCRE experiments

For FLiCRE labeling during nicotine exposure, wildtype mice were used. On day 5 following AAV1/2 injection and fiber implantation in VTA, mice were injected IP with 1.5mg/kg nicotine (delivered in approximately 0.3mL) or saline. Blue light was delivered within 5 min of injecting the mice and placing them in an empty operant box (Coulbourn), using a 470-nm blue LED through a 400-um 0.39 NA patch cable (20 Hz, 10-ms pulse width, ~5 mW power measured at the patch cable, Thorlabs). The stimulation was delivered as 1 min on, 2 min off, for a total duration of 15 min.

For FLiCRE labeling during optogenetic stimulation, Thy1::ChR2 transgenic mice (+ChR2) and wildtype mice (-ChR2) were used. On day 5 following AAV1/2 injection and fiber implantation in mPFC or NAc, mice were placed in an operant box (Coulbourn) where they received 1-10 min of 470-nm blue LED stimulation through a 400-um 0.39 NA patch cable (20 Hz, 10-ms pulse width, ~5 mW power measured at the patch cable, Thorlabs). For the experiment in **Figure S3I**, simultaneous orange and blue light was delivered through the patch cable using a 470/590-nm wavelength division mini cube (Doric, DMC_1×2w_470/590_FC). Orange light was delivered using a 594-nm laser (Cobolt, Mambo). For the -ChR2 wildtype mice in Figure 2, mice were placed under 3% isoflurane to reduce neural activity. However, subsequent experiments did not yield significant differences in labeling when mice were anesthetized versus allowed to freely behave (data not shown), and thus the -ChR2 mice were not anesthetized for data collected in **Figure 6** and **Figure S3**. Animals were placed back in their cage following the blue light delivery. ~18 hrs following the light stimulation, mice were either sacrificed and their brains processed for imaging, electrophysiology, or single-cell RNA sequencing, or underwent the real-time place preference assay.

#### Electrophysiology from FLiCRE-labeled mouse brain slices

On day 5 after AAV1/2 FLiCRE injection and fiber implantation in NAc, a FLiCRE recording was performed in Thy1::ChR2 mice as described above. The following day, the mice were anesthetized deeply with isoflurane and 250 μm coronal slices containing NAc were prepared with a vibratome after intracardiac perfusion with ice-cold artificial cerebrospinal fluid (aCSF) containing (in mM): 126 NaCl, 3 KCl, 1 Na2HPO4, 26.2 NaHCO3, 2.5 CaCl2.2H_2_O, 1.3 MgSO4.7H2O, 11 D-glucose (~7.3 pH, 295-305 mOsm) and oxygenated with 95% O2 / 5% CO_2_. Slices were cut in ice-cold oxygenated sucrose solution containing (in mM): 230 sucrose, 2.5 KCl, 1.25 NaH2PO4, 24 NaHCO3, 0.5 CaCl2.2H2O, 7 MgSO4.7H2O, 11 glucose (~7.3 pH, ~320 mOsm). Following 60 min of recovery in 33°C aCSF, slices were placed in a recording chamber and perfused at 2-4 ml/min with 30°C oxygenated aCSF. Whole cell voltage clamp recordings were conducted using pulled borosilicate glass (G150TF-4, Warner Instruments) patch pipettes (2.5-5.5 MΩ) filled with internal solution containing (in mM): 130 CsMeSO3, 10 HEPES, 0.4 EGTA, 5 TEA-Cl, 7.5 Na2phosophocreatine, 4 MgATP, 0.4 NaGTP, 0.1 spermine, 4 QX-314 Br (7.3 pH, 290-295 mOsm). Labeled (mCherry+) and unlabeled neurons were visualized under a 40× water-immersion objective on a fluorescence microscope (BX51WI; Olympus) with infrared-differential interference contrast and epifluorescence. Input resistance and access resistance were continuously monitored during recordings; experiments were terminated if these changed by >20%.

For EPSC amplitude recordings, ChR2 in presynaptic terminals was stimulated with 470-nm light (5 ms pulses, 5 s apart, 6.5-62.2 μW) through the light path of the microscope using a driver-powered LED (Thorlabs) under computer control. The light intensity of the LED was increased or decreased during experiments to record light power curves and at least two EPSC amplitudes were averaged per power level. A dual lamp house adaptor (Olympus) alternated fluorescence lamp and LED light sources. 100μM picrotoxin (Sigma) was applied during all recordings to block GABAA transmission. Amplitudes were measured from a local baseline 5-10 ms before the light pulse. Slice electrophysiology data were collected with Axograph software.

#### Mouse real-time place preference (RTPP) assay

On day 5 after AAV1/2 FLiCRE injection (TRE-bReaChES-p2a-mCherry reporter) and fiber implantation in NAc, a FLiCRE recording was performed in Thy1::ChR2 and wildtype mice with blue light as described above. Mice were then placed back in their cage. An hour later, mice were placed in a custom-built RTPP chamber (11×2 ft) to determine their baseline preference for each side of the chamber (as described in ref (Kim et al., 2017)). Behavioral tracking was performed using blinded automated software (Biobserve). The following day, mice were stimulated bilaterally with orange light (20 Hz, 10-ms pulse width, ~10 mW power measured at the 400 um 0.39 NA patch cable, Thorlabs) whenever they were on one side of the chamber.

Orange light was delivered using a 594-nm laser (Cobolt, Mambo). Stimulation sides were counterbalanced across mice. Each session lasted 20 min. For Tac1-cre and Calb1-cre mice, animals were allowed to express the AAV5-DIO-ChR2/AAV8-DIO-mCherry for 4-6 weeks, as these serotypes typically take much longer to express (Kim et al., 2017). The animals then underwent the same RTPP assay, but using blue light instead of orange light to stimulate ChR2. Data and statistics analyzed using MATLAB.

#### Confirmation of NAc calcium responses during PFC optogenetic stimulation

Thy1::ChR2 mice were injected with AAV1/2-Synapsin-GCaMP6s in NAc at the same coordinates as described for FLiCRE recording. A 400μM diameter optical fiber was implanted above both the NAc and the PFC. On day 5 after surgery, the bulk NAc GCaMP activity was measured using FIP (Kim et al., 2016). To reduce cross-stimulation of ChR2-expressing axons in NAc, only 5μW of 470-nm light power (measured at the fiber end) was used for NAc photometry recording. A separate 470-nm LED was used to deliver light to the PFC using 10-ms pulses interleaved between photometry excitation pulses. The light power was matched to that used in the real-time place preference assay (~5mW measured at the fiber end).

#### Mouse histology

Following FLiCRE recordings to be analyzed with fluorescence imaging, mice were heavily anesthetized with isoflurane and then perfused with 20 mL of cold phosphate-buffered saline (PBS) followed by 20 mL of cold 4% paraformaldehyde (PFA). The brain was extracted from the skull and incubated in PFA for 24 h, and then transferred to 30% sucrose. After 48 h, the brain was sliced on a vibratome (Leica) in 60-μm sections and mounted on glass slides with coverslips for imaging. For immunohistochemistry of tyrosine hydroxylase (TH), slices were blocked and permeabilized in PBS with 0.3% Triton-X and 5% normal donkey serum. Slices were stained overnight in primary chicken anti-TH (1:500, Aves Lab), and stained for 90 min room temperature in secondary goat anti-chicken alexa-647 (1:1000, ThermoFisher).

#### Single-cell suspensions, barcoded libraries, and RNA sequencing

For single-cell RNA sequencing (scRNA-seq) experiments, a FLiCRE recording was performed in the NAc of Thy1::ChR2 mice as described above for 10 min. The following day, mice were sacrificed and the lateral NAc was micro-dissected to generate a single-cell suspension as previously described (Saunders et al., 2018). The complete protocol is located here, and was followed exactly: www.dropviz.org, under “Data/Single cell suspension protocol from acute adult brain”. We generated 400 μm coronal slices, and microdissected the lateral NAc. The total number of slices collected and pooled across mice for each experimental replicate is listed in **Data File S1**. In total, 3 pooled experimental replicates (including tissue from 5 different mice) were analyzed for sequencing. Tissue was incubated at 34°C for 2 hrs in calcium-free dissociation buffer, supplemented with Protease 23 and Papain. Dissociated cell stocks were eluted in 200 μL prior to cell count estimation.

Following cell count estimation using a Countess II (Thermofisher), barcoded cDNA libraries were prepared (10X Genomics, Chromium Single Cell 3’ v2 kit) at a targeted 800 cells for sequencing. Illumina i7 adapters were incorporated, as per the 10X protocol. cDNA was quantified using an Agilent 2100 Bioanalyzer (Stanford PAN Facility) for all steps except for the final library concentration, which was calculated using a QuBit 4 Fluorometer (ThermoFisher). Libraries were diluted and prepared for sequencing using the NextSeq 500/550 v2.5 High Output Kit, 150 cycles (Illumina). Libraries were sequenced using a NextSeq 550 (Illumina) according to manufacturer specifications. Complete information for each sequencing sample preparation is in **Data File S1**.

#### Analysis of scRNAseq data

Sequencing data were analyzed using the 10X Genomics Cellranger 3.0 software and custom-written Python 3 scripts. Scripts are available at www.github.com/kimck/FLiCRE-analysis-scripts. Briefly, first each experimental replicate was individually processed using cellranger mkfastq and cellranger count. For the genome, we used the 10X provided mouse reference genome, and added the three FLiCRE transcripts (TRE-mCherry, uTEVp, and tTA, which all have unique 3’ ends). We aggregated the three experimental replicates using cellranger aggr (including normalization for sequencing depth).This aggregated dataset was then put through a quality control (QC) process (Zeisel et al., 2018). The following pre-processing was performed:

Remove cells with < 600 UMI counts
Remove cells with < 1.2 UMIs per cell/genes per cell ratio
Remove cells with > 0.4 percent UMIs coming from mitochondrial genes
Remove genes present in < 20 cells

Additionally, we excluded genes from the clustering analysis that were related to sex, neural activity, or floating RNA (Trf, Plp1, Mog, Mobp, Mfge8, Mbp, Hbb-bs, H2-DMb2, Fos, Jun, Junb, Egr1, Xist, Tsix, Eif2s3y, Uty, Kdm5d, Rpl-genes and mt-genes).

QC data were then analyzed for secondary dimensionality reduction and clustering using cellranger reanalyze (default parameters). QC data were also initially explored using Seurat (Stuart et al., 2019) and Scanpy (Wolf et al., 2018), and yielded similar clustering results. The cellranger clustering scripts were used for simplicity.

For clustering of the tsne embeddings, k-means clustering (10 clusters) was used. For the initial clustering of all cell types, outliers (likely representative of doublets) were removed by setting a threshold of 0.5 euclidean distance from the median center of mass of each cluster. This distance calculation was scaled by the size of each cluster in the x and y dimension. UMI counts were normalized by a scale factor representing the total UMI counts per cell/median UMI counts per cell. Differentially expressed genes across clusters were identified by the cellranger reanalyze algorithm, which tests whether the in-cluster mean gene expression differs from the out-of-cluster mean. <u>Differentially expressed genes were filtered to those with an average UMI count > 1</u>. Cell-types were identified by cross-referencing the top differentially-expressed genes with previously published datasets (aggregated on the Single Cell Portal from the Broad Institute, and from refs (Saunders et al., 2018; Zeisel et al., 2018)).

Clusters identified as neuronal sub-types were then re-clustered using cellranger reanalyze (num_pca_genes=100, num_principal_comps=20, tsne_perplexity=30, tsne_input_pcs=10). In this case, no outliers were removed from clusters. One small cluster that was distributed widely across tsne space was removed as it likely belonged to contaminants/doublets. In addition to the genes excluded from the clustering analysis described above, we also excluded the three FLiCRE transcripts. Data were re-analyzed as before.

To control for different levels of tTA and uTEVp transcripts expression among neurons that could possibly account for mCherry expression levels, we took two approaches. First, we thresholded neurons to identify those that had normalized uTEVp or tTA UMI counts > 2 S.D. of the entire neuron distribution. We then re-clustered these neurons using cellranger reanalyze (num_pca_genes=100, num_principal_comps=20, tsne_perplexity=30, tsne_input_pcs=10). The three FLiCRE transcripts were not excluded from this clustering analysis. The uTEVp and tTA UMI counts were not significantly different across the clusters identified using this method.

Second, we calculated an mCherry enrichment score among neurons that expressed uTEVp or tTA (FLiCRE+). This was calculated as previously described (Moffitt et al., 2018), by taking the percentage of FLiCRE+ neurons in each cluster that were also mCherry+ and dividing that by the total number of mCherry+/FLiCRE+ neurons across all experimental replicates. This enrichment score was calculated individually for each experimental replicate. mCherry+ neurons were defined as those having normalized mCherry UMI counts > 1. FLiCRE+ neurons were defined as those having a normalized uTEVp or tTA UMI counts > 1.

To identify genes significantly enriched in mCherry+/FLiCRE+ neurons, for each gene we compared the true mean normalized UMI count to 10,000 randomly calculated mean normalized UMI counts (where the mCherry+ neurons were randomly selected among FLiCRE+ neurons). Genes that had true UMI counts below the 5^th^ or above the 95^th^ percentile of the shuffled distribution were identified as being significantly down-or up-regulated, respectively.

All scRNA-seq data analysis, statistics, and figure generation were performed using open-source software in Python 3: Matplotlib (Hunter, 2007), SciPy (Jones et al., 2001), Numpy (Oliphant, 2006), Pandas (McKinney, 2010), Seaborn (https://seaborn.pydata.org/), and Statsmodels (Seabold and Perktold, 2010).

#### Luciferin-gated FLiCRE experiment

Either cortical or hippocampal neurons were prepared and infected with FLiCRE viruses as described above. For GCaMP5G imaging of basal versus high calcium activity during electric field stimulation in **Figure 7C** and **7F**, 100μL of crude supernatant AAV1/2-Synapsin-GCaMP5G virus was added to the wells. Imaging was performed using SlideBook, and the field stimulation was delivered as described above (50 pulses delivered at 20Hz every 10 s, 1-ms pulse width, 48 mA). For FLiCRE labeling, neurons were exposed to blue light for 15 min (2s on/4s off) following either a 50% media change (induces high neural activity) or left untreated. For luciferin-gated FLiCRE labeling, neurons were exposed to a final concentration of ~10μM furimazine (1:100 dilution of Promega Nano-Glo Live Assay substrate) following either a 50% media change or left untreated. The furimazine was not washed out following the 15-min experiment.

#### Considerations for *in vivo* FLiCRE labeling

All of the FLiCRE experiments performed here were using homemade concentrated AAV1/2 viruses, as described in (Konermann et al., 2013), and thus only required 5 days of expression. If using AAV8 or AAVDJ, we recommend following the incubation guidelines for those serotypes. While we did not observe any toxicity while expressing FLiCRE using our serotypes over ~1 week of expression, we did not assess potential toxicity effects of long-term expression of FLiCRE. To ensure the calcium-dependency of FLiCRE labeling, it is crucial that the components are not over-expressed – particularly of the TEV protease. For example, if the levels of the protease component are highly expressed, the TEVp/TEVcs cleavage can occur even in the absence of a calcium-induced protein-protein interaction. As such, when expressing the three FLiCRE components *in vivo,* it is critical to ensure the ratio of the viruses is such that the TEVp component is lower than the other two components. We suggest using the ratios reported in **Table S2** as a starting point, and diluting the entire volume mixture with PBS if background FLiCRE expression is observed in the absence of blue light and a strong behavioral stimulus. Our characterization *in vivo* suggests that FLiCRE can detect at least a 5-fold increase in neural activity over background when delivering 20 Hz optogenetic stimulation; however, in brain regions with high, persistent background firing rates of 20Hz or more, FLiCRE may not be suitable. Once an appropriate viral dilution has been established using both a negative and positive control *in vivo* in the brain region of interest, we then recommend using that same dilution for an *in vivo* FLiCRE labeling during the user’s desired stimulus. Furthermore, the longest duration of *in vivo* FLiCRE labeling performed here was for 15 min, while delivering 20Hz blue light pulses for 1 min on, and 2 min off to reduce the overall blue light exposure and background labeling. We suggest that if the user desires to record neural activity over long periods of time (>15 min), that they stagger the blue light as we did, rather than delivering blue light continuously during the behavior, in order to reduce background labeling. Finally, we suggest waiting ~18-24 hrs following the FLiCRE recording prior to sacrificing mice to look for reporter gene expression (or for optogenetic experiments). Waiting significantly longer could increase background expression.

### QUANTIFICATION AND STATISTICAL ANALYSIS

The exact statistical test, n-value and description, precision measures, and definition of center is listed in the figure legend of every data panel. Statistical analysis was performed using the statistics toolbox in MATLAB R2017a (Mathworks) and Python 3 (packages listed in section “Analysis of scRNAseq data”). Significance was generally defined as a P-value less than 0.05 for the defined statistical test; except when an FDR adjusted P-value was determined (noted in figure legend). No statistical methods were used to pre-determine sample size or fit with assumptions of statistical tests. No subjects were excluded from the study. Cells not passing the basic quality control metrics defined in “Single-cell suspensions, barcoded libraries, and RNA sequencing” were omitted from the study.

### SUPPLEMENTAL MATERIALS

**Data File S1. Related to Figure 4. Information about sequencing libraries.**

Excel sheet containing parameters for single-cell suspensions, library generation, and sequencing.

