## Supplemental Information for "A molecular calcium integrator reveals a striatal cell-type driving aversion"

**Table S1. Related to STAR Methods. Plasmids used or generated in this study.**

| **Name** | **Description** | **Vector-Promoter** | **Access** |
| --- | --- | --- | --- |
| CD4-MKII-eLOV-TEVcs(ENLYFQ/M)- Gal4 | Transcription factor (TF) component of FLARE | AAV-CMV | Addgene #92213 |
| GFP-CaM-TEVp | Protease component of FLARE fused to GFP | AAV-CMV | Available upon request |
| CD4-MKII-iLID-TEVcs(ENLYFQ/M)- Gal4 | TF component of FLARE with iLID variant | AAV-CMV | Available upon request |
| CD4-MKII-hLOV2-TEVcs(ENLYFQ/M)- Gal4 | TF component of FLARE with hLOV2 variant | AAV-CMV | Available upon request |
| UAS:mCherry | FLiCRE reporter gene for HEK293T expression | AAV-UAS | Addgene #135457 |
| CD4-MKII-hLOV1-TEVcs(ENLYFQ/M)- Gal4 | FLiCRE TF for HEK293T expression | AAV-CMV | Addgene #163026 |
| CD4-MKII-f-hLOV1-TEVcs(ENLYFQ/M)- Gal4 | FLiCRE TF with fast kinetics for HEK293T expression | AAV-CMV | Addgene #163027 |
| GFP-CaM-uTEVp | FLiCRE protease for HEK293T expression | AAV-CMV | Addgene # 163028 |
| Nrxn3b-Nav1.6-MKII-hLOV1-TEVcs(ENLYFQ/M)-tTA-VP16 | FLiCRE TF for neuronal expression | AAV-Synapsin | Addgene #163029 |
| Nrxn3b-Nav1.6-MKII-f-hLOV1-TEVcs(ENLYFQ/M)-tTA-VP16 | FLiCRE TF with fast kinetics for neuronal expression | AAV-Synapsin | Addgene # 163031 |
| GFP-CaM-uTEVp | FLiCRE protease for neuronal expression | AAV-Synapsin | Addgene #163032 |
| mtagBFP2-CaM-uTEVp | FLiCRE protease for neuronal expression | AAV-Synapsin | Addgene #163033 |
| GFP-CaM-NanoLuc-15aa-uTEVp | FLiCRE protease with luciferin-gating for neuronal expression | AAV-Synapsin | Addgene #163034 |
| TRE:mCherry | FLiCRE reporter gene for neuronal expression | AAV-TRE | Addgene #92202 |
| TRE:GFP | FLiCRE reporter gene for neuronal expression | AAV-TRE | Addgene #163036 |
| TRE:mCherry-p2a- bReaChES | FLiCRE reporter gene for neuronal expression | AAV-TRE | Addgene #163037 |
| bReaChES-p2a-mCherry | Excitatory opsin | AAV-Synapsin | Available upon request |
| GCaMP5G | Calcium indicator | AAV-Synapsin | Available upon request |
| M13-TEV-C-p2a-eGFP | One of the components used in Cal-Light | AAV-Synapsin | Available upon request |
| [TM-CaM-NES-TEV-N-AsLOV2-TEVseq-tTA](https://www.addgene.org/92392/) | Membrane-anchored TF component used in Cal-Light | AAV-Synapsin | Addgene #92392 |
| MBP-LOVwt | Bacterial expression of LOV domains fusions | YFJ16-LacZ | Available upon request |
| MBP-eLOV | Bacterial expression of LOV domains fusions | YFJ16-LacZ | Available upon request |
| MBP-hLOV1 | Bacterial expression of LOV domains fusions | YFJ16-LacZ | Available upon request |
| MBP- f-hLOV1 | Bacterial expression of LOV domains fusions | YFJ16-LacZ | Available upon request |

**Table S2. Related to STAR Methods. Virus volumes and concentrations used for experiments.**

| **Figures** | **Virus** | **Source** | **Stock titer** |
| --- | --- | --- | --- |
| 2, S1C-S1E, S2A | Per 24-well:   - 100µL AAV1/2-Synapsin-Nrxn3b-MKII-hLOV1-TEVcs-tTA - 100µL AAV1/2-Synapsin-GFP-CaM-uTEVp - 100µL AAV1/2-TRE-mCherry | In-house | N/A – crude supernatant |
| 3A-3C | Per injection:   - 0.1µL AAV1/2-Synapsin-Nrxn3b-MKII-hLOV1-TEVcs-tTA - 0.025µL AAV1/2-Synapsin-GFP-CaM-uTEVp - 0.075µL AAV1/2-TRE-mCherry - 0.3µL PBS | In-house | 1.38e9 vg/µL  2.58e9 vg/µL  1.38e9 vg/µL |
| 3D-3L, 4 | Per injection:   - 0.4µL AAV1/2-Synapsin-Nrxn3b-MKII-hLOV1-TEVcs-tTA - 0.1µL AAV1/2-Synapsin-GFP-CaM-uTEVp - 0.3µL AAV1/2-TRE-mCherry - 0.2µL PBS | In-house | 1.38e9 vg/µL  2.58e9 vg/µL  1.38e9 vg/µL |
| 6C-6D | Per injection:   - 0.4µL AAV1/2-Synapsin-Nrxn3b-MKII-hLOV1-TEVcs-tTA - 0.1µL AAV1/2-Synapsin-GFP-CaM-uTEVp - 0.3µL AAV1/2-TRE-mCherry-p2a-bReaChES - 0.2µL PBS | In-house | 1.38e9 vg/µL  2.58e9 vg/µL  2.0e9 vg/µL |
| 7C,7F | Per 24-well:   - 100µL AAV1/2-Synapsin-GCaMP5G | In-house | N/A – crude supernatant |
| 7D,7G | Per 24-well:   - 100µL AAV1/2-Synapsin-Nrxn3b-MKII-hLOV1-TEVcs-tTA - 100µL AAV1/2-Synapsin-GFP-CaM-NanoLuc-15aa-uTEVp - 100µL AAV1/2-TRE-mCherry | In-house | N/A – crude supernatant |
| S2B, S2E-S2H | Per 24-well:   - 100µL AAV1/2-Synapsin-M13-TEV-C-p2a-eGPF - 100µL AAV1/2-Synapsin- Nrxn3b-CaM-NES-TEV-N-AsLOV2-TEVcs-tTA - 100µL AAV1/2-TRE-mCherry | In-house | N/A – crude supernatant |
| S3C | Per injection:   - 0.5µL AAV1/2-Synapsin-Nrxn3b-MKII-f-hLOV1-TEVcs-tTA - 0.04µL AAV1/2-Synapsin-BFP-CaM-uTEVp - 0.2µL AAV1/2-TRE-mCherry - 0.2µL PBS | In-house | 8.60e8 vg/µL  9.70e9 vg/µL  1.38e9 vg/µL |
| S3I | Per injection:   - 0.4µL AAV1/2-Synapsin-Nrxn3b-MKII-hLOV1-TEVcs-tTA - 0.05µL AAV1/2-Synapsin-BFP-CaM-uTEVp - 0.3µL AAV1/2-bReaChES-p2a-mCherry - 0.02µL AAV1/2-TRE-eGFP - 0.2µL PBS | In-house | 1.38e9 vg/µL  9.70e9 vg/µL  2.00e9 vg/µL  2.5e10 vg/µL |
| 6G-6J | Per injection:   - 1µL AAV5-EF1a-DIO-ChR2-eYFP, **or** - 1µL AAV8-EF1a-DIO-mCherry | UNC core  Stanford core | 5.0e12 vg/mL  1.7e12 vg/mL |
| S6F | Per injection:   - 1µL AAV1/2-Synapsin-GCaMP5G | In-house | 6.5e9 vg/µL |

**Table S3. Related to Figure 7. Summary of SNRs reported for activity-dependent labeling technologies.**

|  | | **c-fos-tTA mice** | **TRAP2 mice** | **Cal-Light** | **FLARE** | **CaMPARI2** | **FLiCRE** |
| --- | --- | --- | --- | --- | --- | --- | --- |
| ***In vitro*** | **~SNR** | **Not reported** | **Not reported** | **4x** | **11x** | **23x** | **66x** |
|  | **+/- activity in cultured neurons** |  |  | **+/- 4 hrs bicuculline** | **+/- 8 min 20Hz field stimulation** | **+/- 2 sec 80Hz field stimulation** | **+/- 10 min 20Hz field stimulation** |
| ***In vivo*** | **~SNR** | **2x** | **2x or 9x** | **3x** | **Not reported** | **6x** | **4x, 5x, or 6x** |
|  | **+/- activity (region)** | **+/- 3 fear conditioning sessions over 9 hrs (mouse DG)** | **+/- 1 fear conditioning session (mouse PFC)**  **+/- 6 hrs water deprivation (mouse MnPO)** | **+/- 6 days of 45 min leverpress sessions (mouse M1)** |  | **+/- 30 sec swimming  (larval zebrafish forebrain)** | **+/- 15 min nicotine (mouse VTA)**  **+/- 1 min direct stimulation (mouse PFC)**  **+/- 10 min of input stimulation (mouse NAc)** |
| **Reference** | | **Liu et al., *Nature* 484, 381-385 (2012), Fig 2H.** | **Denardo et al., *Nature Neuro* 22, 460-469 (2019), Fig S4C; Allen et al., *Science* 357, 1149-1155 (2017), Fig 1F.** | **Lee et al., *Nature Biotech* 35, 858-863 (2017), Fig 1D, Fig 2I.** | **Wang et al., *Nature Biotech* 35, 864-871 (2017), Fig 3E.** | **Moeyaert et al., *Nature Comm* 9, 4440 (2018), Fig 1D, Fig 3B.** | **This manuscript, Fig S2E, Fig 3C, S3E, Fig 3I.** |
